# Toroidal Search Algorithm: A Topology-Inspired Metaheuristic with Applications to ODE Parameterization in Mathematical Oncology

**DOI:** 10.64898/2026.03.05.709766

**Authors:** Changin Oh, Kathleen P. Wilkie

## Abstract

We present the Toroidal Search Algorithm (TSA), a novel population-based metaheuristic optimization method inspired by the topology of a torus. Conventional metaheuristics frequently suffer from boundary stagnation, a phenomenon that severely degrades performance in bounded and high-dimensional search spaces. TSA addresses this limitation by embedding the search domain into a toroidal geometry, thereby eliminating artificial boundaries and enabling continuous cyclic exploration. Beyond boundary handling, TSA uses winding numbers to capture the history of agent movement across periodic dimensions, which are exploited to adaptively refine local search. A modified sigmoid control function regulates the transition between global and local search. Performance of TSA is evaluated on a collection of unimodal and multimodal benchmark functions at various dimensions. It consistently outperforms established metaheuristics. Notably, TSA demonstrates exceptional robustness to increasing dimensionality, maintaining fast convergence and low variance where competing methods deteriorate. To assess real-world applicability, we apply TSA to an inverse problem from mathematical oncology. With both synthetic and clinical data, TSA reliably recovers physiologically plausible parameters with greater stability and predictive accuracy than competing algorithms. These results demonstrate that TSA is a powerful and robust tool for large-scale global optimization in computational modelling applications.

**Striking Image:** 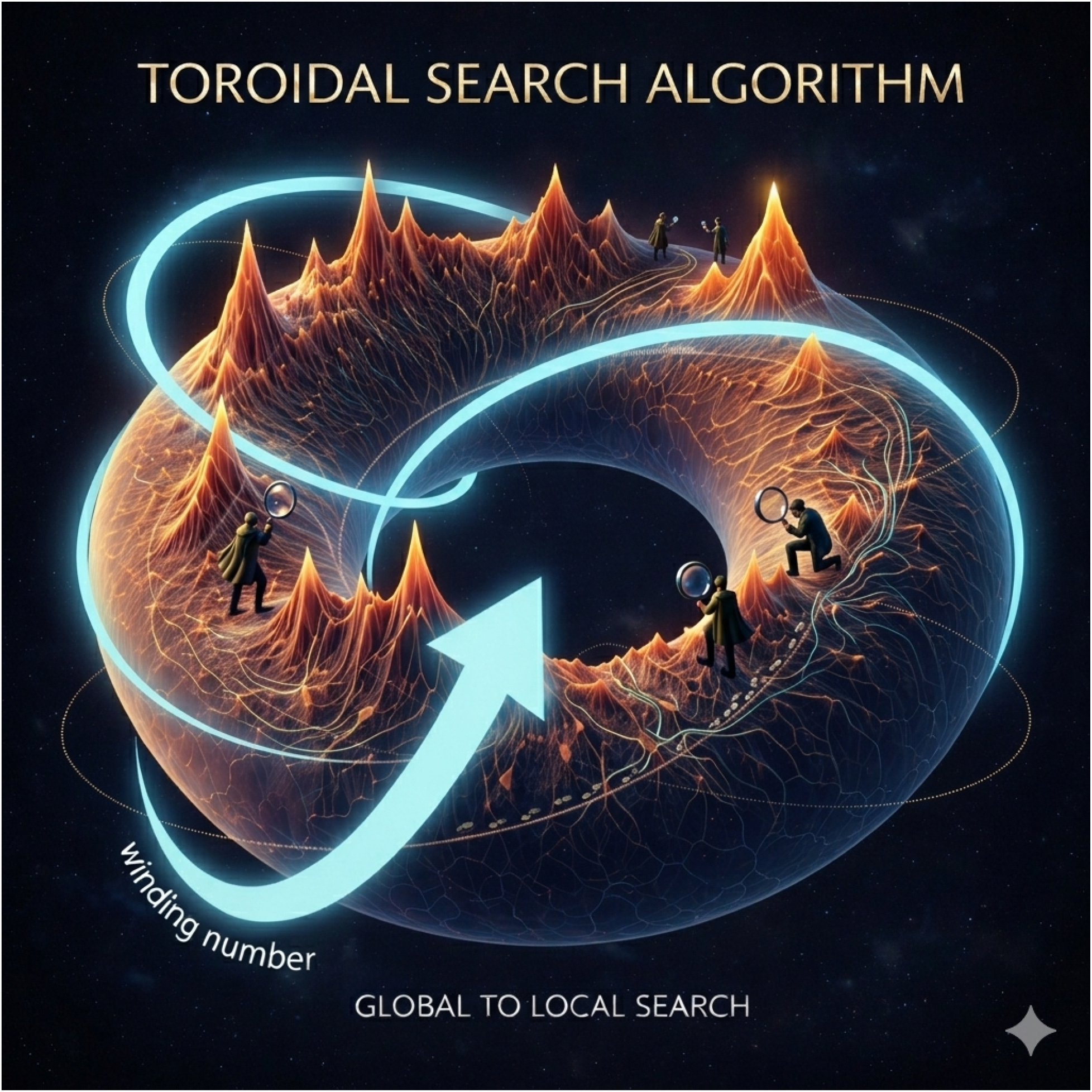

Image generated with Google Gemini.

## 1 Introduction

Optimization is the process of finding the “best” solution to a problem from a set of possible alternatives, and because many real-world systems are complex, nonlinear, and high-dimensional, optimization provides the mathematical and computational tools needed to make effective decisions. Optimization, a key component of uncertainty quantification, involves maximizing or minimizing an objective function subject to given constraints. Problems of this type appear in almost every field of science, engineering, medicine, and industry, ranging from the design of efficient transportation networks [1] to the training of machine learning models [2].

In recent years, the importance of optimization has grown significantly. The emergence of big data and artificial intelligence has created a demand for algorithms that can handle large problem sizes and extract meaningful patterns in reasonable time [3]. In addition, global challenges such as climate change [4], sustainable resource management [5], drug discovery [6], and personalized medicine [7], increasingly require solving difficult optimization problems that cannot be addressed by traditional deterministic methods. Metaheuristic algorithms, in particular, have attracted attention because of their flexibility and ability to approximate global solutions in complex search spaces.

Metaheuristic algorithms are often inspired by natural and physical phenomena, leading to diverse approaches such as Particle Swarm Optimization (PSO) [8], Differential Evolution (DE) [9], Firefly Algorithm (FA) [10], Artificial Bee Colony (ABC) [11], and Teaching-Learning-Based Optimization (TLBO) [12]. Each of these methods has been successfully applied to different classes of optimization problems. However, the No Free Lunch (NFL) theorem for optimization [13] states that, when performance is averaged over all possible problem classes, all algorithms are equivalent. This means that an algorithm that performs well on certain problem classes will inevitably perform worse on others. Therefore, it is essential to design new optimization methods that are based on the specific characteristics of the problems being addressed.

Boundary handling is an important consideration in the design and evaluation of metaheuristic algorithms. Many optimization problems have strict upper and lower bounds on decision variables, and the way an algorithm handles these boundaries can significantly influence its performance. Inappropriate handling can distort the search trajectory, limit exploration, or disrupt convergence behaviour [14, 15, 16].

Simple boundary handling strategies such as absorbing, reflection, or random reinitialization are commonly adopted because they are easy to implement and add little computational cost [14]. In absorbing strategies, solutions that move outside the boundary are forced back to the boundary. Reflection sends solutions back into the feasible region by mirroring them across the boundary. And random reinitialization discards out-of-bounds solutions and replaces them with new ones sampled uniformly within the search domain. However, these strategies can lead to a serious limitation known as boundary stagnation [14, 17, 18], where the search process gets trapped near the edges of the domain. This causes premature convergence to suboptimal solutions and degrades performance, especially in high-dimensional problems where boundary violation occurs much more frequently [19].

Addressing boundary handling has the potential to significantly improve the robustness and efficiency of metaheuristic optimization, particularly in modern applications where problems are highly sensitive to search dynamics. Among the various boundary handling methods, the wrapping strategy offers a different perspective. In this approach, the search space is treated as if it were wrapped onto a torus, so that solutions exiting one boundary re-enter from the opposite side [20]. This eliminates the presence of hard edges and allows continuity of the search process, thus reducing the risk of boundary stagnation. Despite its conceptual elegance, previous studies have shown that many existing metaheuristic algorithms do not perform well when wrapping is applied as a boundary handling mechanism [21], since their search operators were not originally designed for toroidal geometry. This observation motivates the development of a new algorithm that incorporates the toroidal structure at its core.

To this end, we propose the Toroidal Search Algorithm (TSA), a novel metaheuristic that integrates wrapping boundary handling as a fundamental design principle. TSA takes advantage of the toroidal property not only for boundary constraint handling but also as a core search strategy. By mapping the search space onto a torus, TSA enables cyclic exploration, which reduces boundary stagnation and provides multiple paths for search agents to approach the optimal solution, thereby enhancing global search. Furthermore, TSA introduces winding numbers to track cyclic movements, which refine local search by providing information about repeated traversals of the toroidal space. Finally, the integration of a sigmoid function dynamically balances exploration and exploitation, allowing the transition from global search to local search.

Below, we introduce TSA, detailing its theoretical basis, algorithmic structure, and key features. We evaluate the performance of TSA through extensive benchmarking on standard optimization problems, and compare it with established algorithms such as PSO, DE, FA, ABC, and TLBO. The results show that TSA achieves competitive or improved performance, highlighting its robustness and efficiency, even in complex, high-dimensional search spaces.

The remainder of this paper is organized as follows. Section 2 provides the necessary mathematical concepts underlying TSA. Section 3 describes TSA, including its design principles and search mechanisms. Section 4 provides the experimental setup and benchmark functions. Section 5 reports and discusses all benchmarking results. Section 6 demonstrates the real-world application of TSA to an ODE parameterization problem from mathematical oncology. Section 7 concludes the paper with final remarks and directions for future work.

## 2 Mathematical Background

### 2.1 Torus

A torus is a topological structure that extends the idea of periodicity to higher dimensions. It is defined as the Cartesian product of circles, representing a space with periodic boundary conditions. An *n*-dimensional torus (or simply *n*-torus) is given by

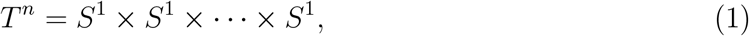

where *S*^1^ denotes the unit circle. Equivalently, an *n*-torus can be constructed by identifying the opposite faces of an *n*-dimensional hypercube.

The simplest case, the 1-torus or *T* ^1^, is just the unit circle, which can be though of as a line segment with its endpoints glued together. The 2-torus or *T* ^2^ is often visualized as a donut-shaped surface and can be constructed by gluing together the opposing sides of a square (Figure 1). Higher-dimensional tori such as *T* ^3^ and beyond cannot be visualized in three-dimensional space but preserve the same periodic structure.

**Figure 1.**
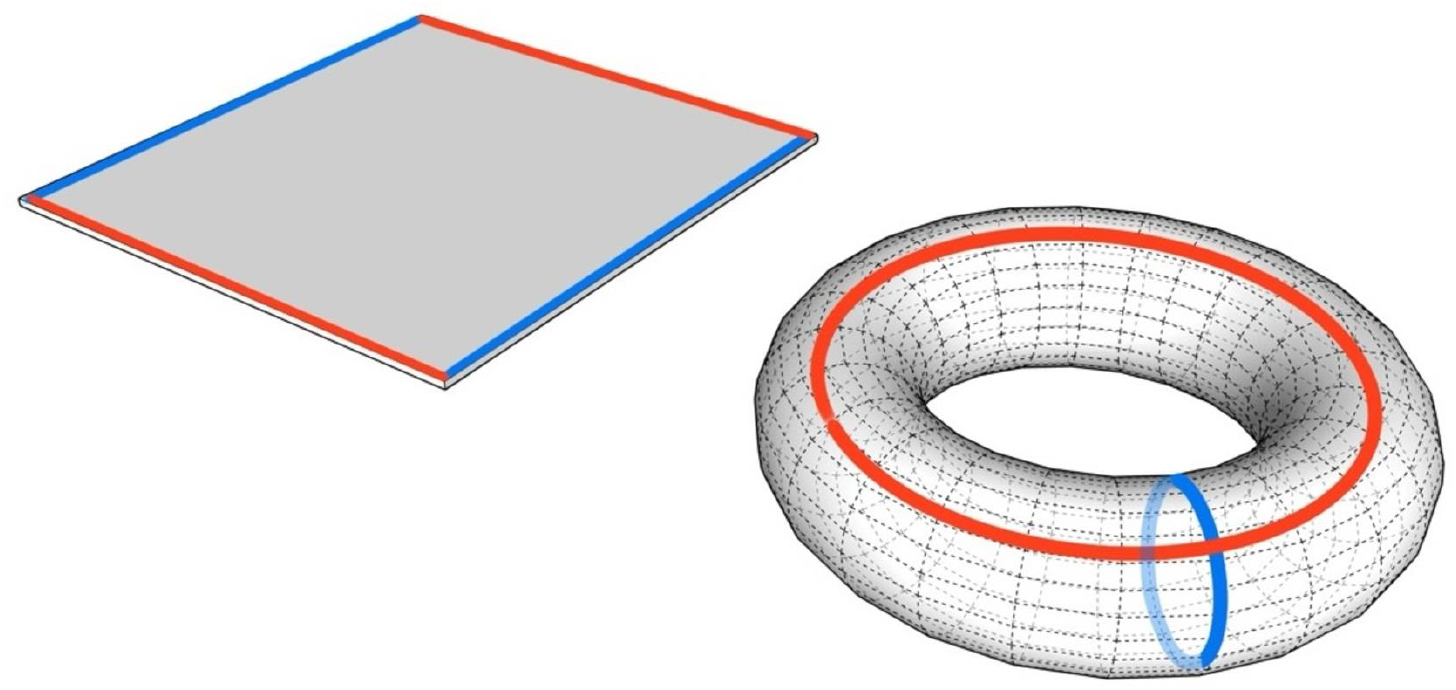
Illustration of a 2-torus or *T* ^2^. The donut-shaped surface (right) is obtained by gluing together the opposite sides of a rectangle (left).

Due to the periodic structure of a torus, the representation of any point is not unique. Each point is repeated after a fixed interval, so that moving indefinitely in one direction will repeatedly return to the starting point (Figure 2). This property establishes periodic boundary conditions: leaving through one face of the hypercube corresponds to immediate re-entry from the opposite face.

**Figure 2.**
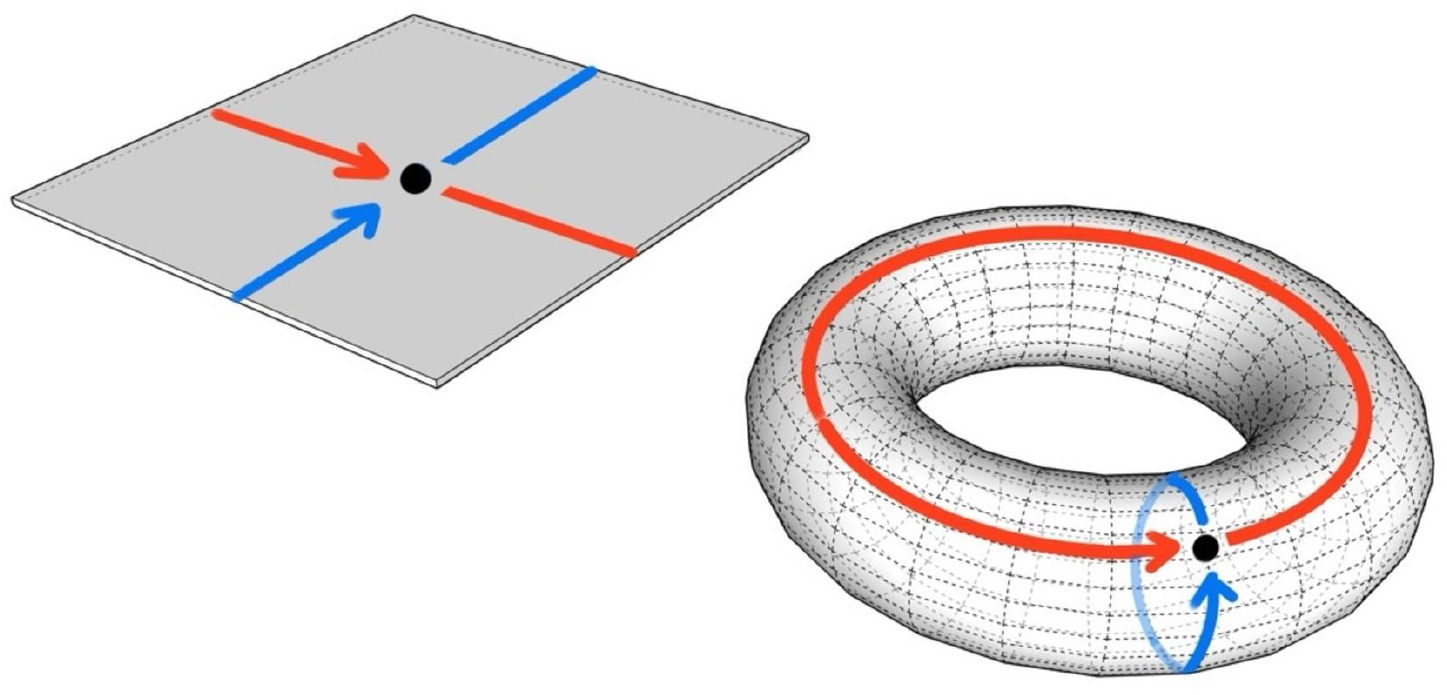
Illustration of periodic boundary conditions on a 2-torus. Exiting one side of the rectangle results in immediate re-entry from the opposite side (left). The corresponding movements are shown on the 2-torus (right).

Mathematically, periodic boundary conditions in toroidal spaces can be expressed using real-valued modular arithmetic, which allows continuity by wrapping values within a predefined range. For simplicity, consider the 2-torus. Let [*a, b*] × [*c, d*] be a rectangle of width *w* = *b* − *a* and height *h* = *d* − *c*. Define the equivalence relation ∼ by

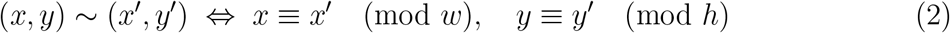

for any *x, x*^*′*^ ∈ [*a, b*] and *y, y*^*′*^ ∈ [*c, d*]. This relation states that points (*x, y*) and (*x*^*′*^, *y*^*′*^) are considered equivalent if their coordinates differ by integer multiples of the width and height of the rectangle. The quotient space [*a, b*] × [*c, d*] */* ∼ therefore identifies opposite sides of the rectangle, forming a 2-torus. In this representation, a point that exits through one boundary immediately re-enters from the opposite boundary, yielding an uninterrupted, continuous search domain. This construction extends naturally to higher dimensions, where the *n*-torus is obtained by imposing periodicity in each coordinate direction.

### 2.2 Winding Numbers

The winding number is a concept used to describe how motion behaves in spaces with periodic structure. Informally, it counts how many times a path loops around a space before returning to a previously visited location.

A useful way to understand winding numbers comes from topology, where paths are classified according to how they can be continuously deformed. Two paths are said to belong to the same homotopy class if one can be smoothly deformed into the other without cutting the path, even if their detailed shapes are different.

On a circle, this idea leads to a particularly clear interpretation. Any closed path on a circle can be continuously deformed into a path that winds around the circle a certain number of times. Paths that wind different numbers of times cannot be continuously deformed into one another without breaking the path or leaving the circle, and therefore belong to different homotopy classes. As a result, the homotopy classes of closed paths on a circle are indexed by integers. In formal terms, the fundamental group of *S*^1^ is isomorphic to the (additive) group of integers,

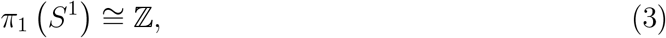

where each integer records the number of net revolutions around the circle with the sign indicating direction. This integer is called the winding number. Positive integers correspond to counterclockwise motion and negative integers to clockwise motion. This interpretation is illustrated in Figure 3, which shows representative closed paths with winding numbers of −1, −2, +1, and +2.

**Figure 3.**
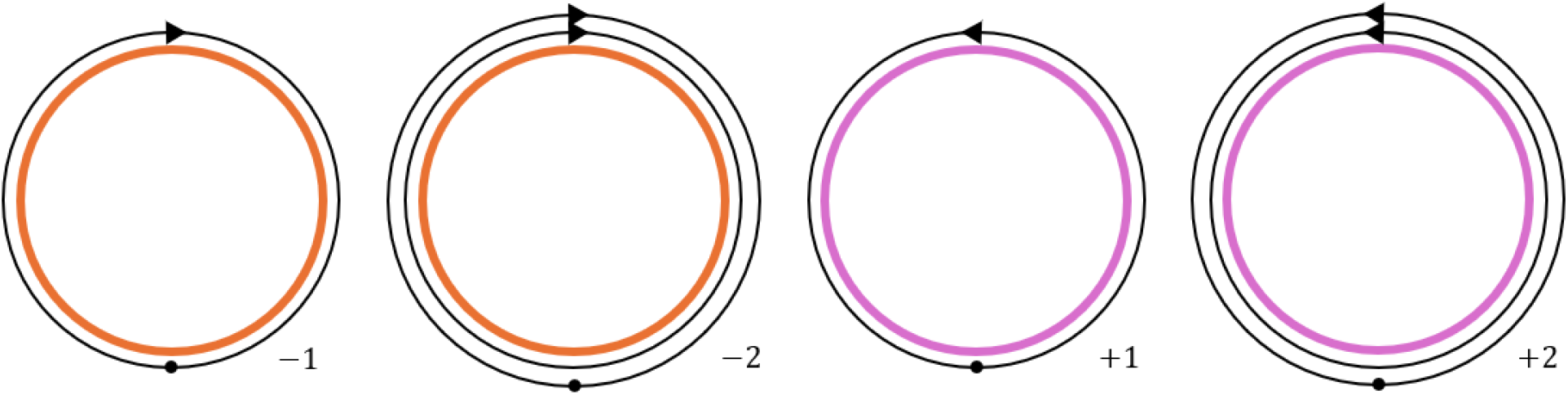
Illustration of winding numbers on *S*^1^.

This idea generalizes naturally to higher-dimensional periodic spaces such as a torus. An *n*-dimensional torus can be viewed as a space in which each coordinate direction is periodic. From a topological perspective, this structure implies that loops on the torus can wrap independently around each coordinate direction. In the case of the two-dimensional torus *T* ^2^, there are two fundamental types of loops: one that winds around the torus along the longitudinal direction and another that winds along the meridional direction. These two independent circular directions are illustrated in Figure 4, where each loop corresponds to *S*^1^ embedded in the torus. Consequently, the homotopy classes of loops on an *n*-dimensional torus can be described by *n* winding numbers. Mathematically, the fundamental group of the *n*-torus satisfies

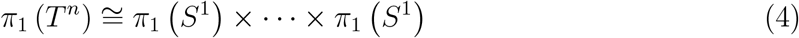

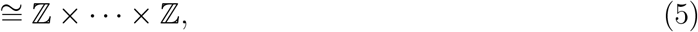

meaning that each loop can be described by an *n*-tuple of integers (or an integer vector in Z^*n*^), where each component counts the number of times the loop winds around the torus in a particular dimension.

**Figure 4.**
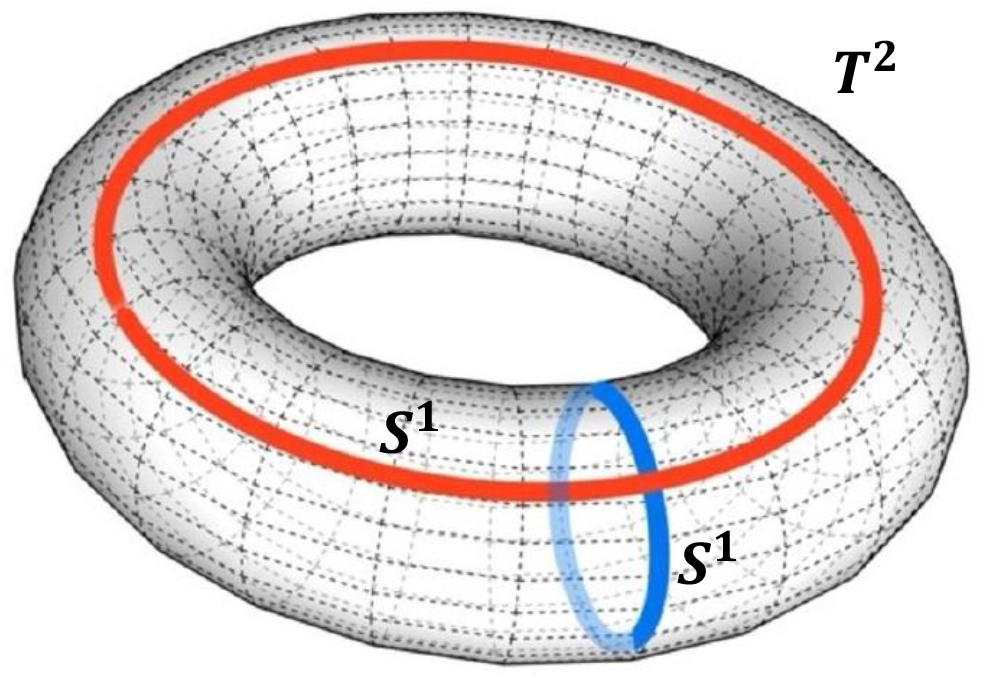
Illustration of *T* ^2^ and its two independent circular directions. The red and blue curves represent two distinct fundamental groups of *S*^1^ corresponding to loops that wind around the torus.

### 2.3 Sigmoid Function

The sigmoid function is a mathematical function that produces an S-shaped curve, mapping real numbers to the bounded interval (0,1). It is widely used in machine learning [22] and control systems [23] because of its smoothness, differentiability, and ability to regulate gradual transitions between states. The general form is

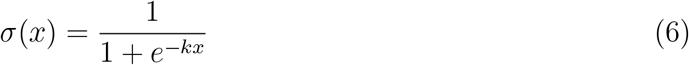

where *k* ≥ 0 controls the steepness of the curve. The function is strictly increasing, symmetric around zero where it evaluates to 0.5, and smoothly transitions from 0 to 1 as *x* increases. These properties make it useful for scaling values and dynamically adjusting behaviour in algorithms.

In the context of metaheuristic algorithms that begin with a global search and transition to a local search, a modified sigmoid function can be used to shift between the two phases. Specifically,

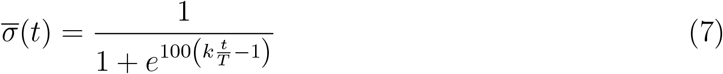

where *t* is the current iteration, *T* is the maximum number of iterations, and *k* determines the transition point. The scaling factor of 100 in the exponent causes an abrupt shift rather than a gradual one. In early iterations, the large negative exponent keeps 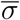 close to 1, maintaining a global search. In later iterations with large *t*, however, the function rapidly approaches 0, transitioning the algorithm to a local search. The control parameter *k* determines the iteration-timing of this switch. As shown in Figure 5, for *k* > 2, global search ends relatively early, while *k* < 2 prolongs exploration before switching to local search. Setting *k* = 2 yields a balanced transition, allocating equal effort to both global and local searches.

**Figure 5.**
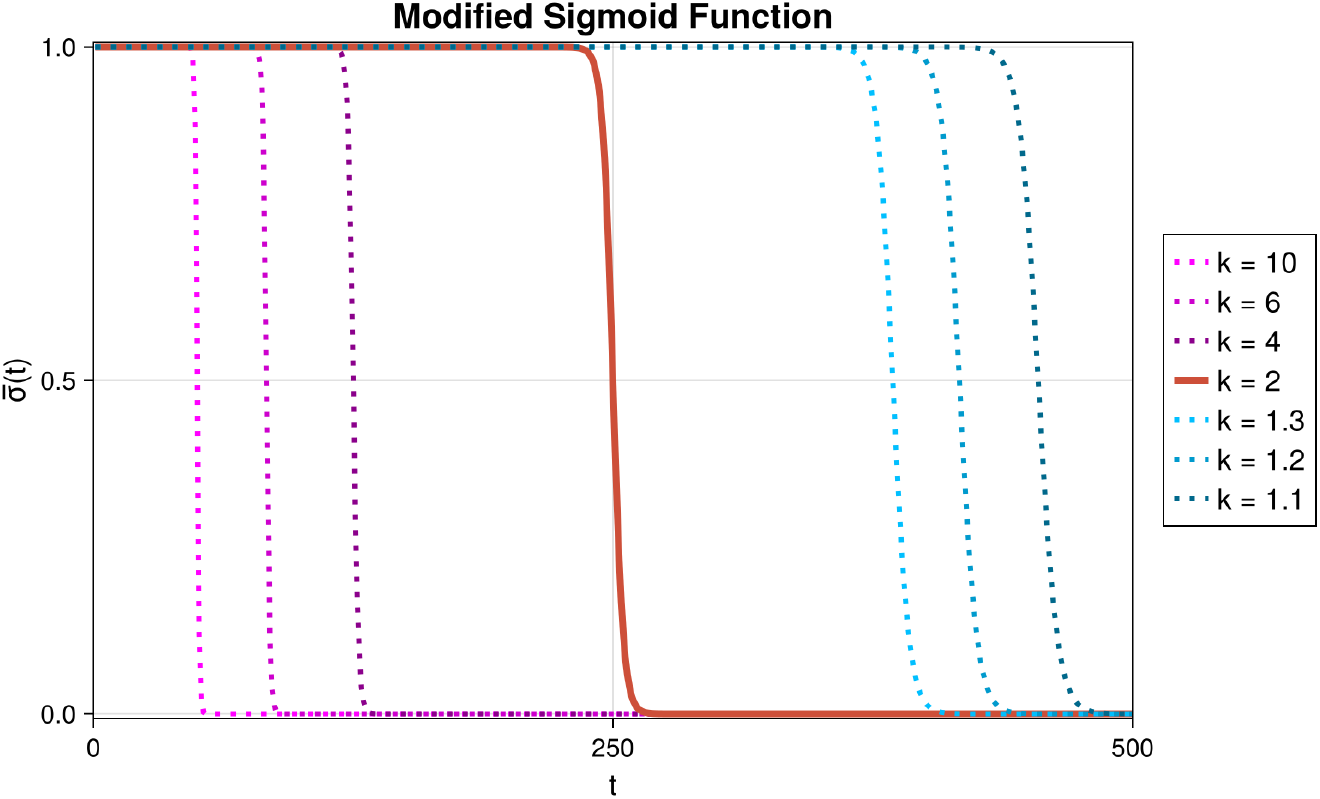
Illustration of the modified sigmoid function used in TSA (*T* = 500). When *k >* 2, the global search ends quickly, whereas with *k <* 2 the global search lasts longer. When *k* = 2, equal effort is allocated to both global and local searches.

## 3 Toroidal Search Algorithm

TSA is designed as a metaheuristic optimization algorithm that integrates toroidal geometry, winding numbers, and a modified sigmoid function into its search process. The update mechanism of TSA consists of two components: a global search term and a local search term. Mapping the search space onto a torus eliminates boundary stagnation and enables cyclic global search while winding numbers refine the local search by scaling agent movements to smaller steps. The modified sigmoid function dynamically balances these two phases, which allows TSA to perform global search during the early iterations and transition to local search in later iterations.

### 3.1 Initialization

TSA begins with a population of *N*_*p*_ candidate agents, each denoted as

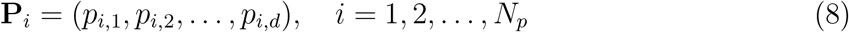

for a *d*-dimensional search space. Each agent is initialized uniformly at random within the lower and upper bounds of the search domain:

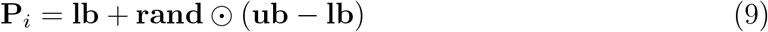

where **rand** is a vector of random values sampled from the uniform distribution over [0, 1], and **lb** = (*l*_1_, *l*_2_, …, *l*_*d*_) and **ub** = (*u*_1_, *u*_2_, …, *u*_*d*_) are the lower and upper bounds of the *d*-dimensional search space. Operator ⊙ represents the Hadamard product, i.e., element-wise multiplication. The fitness of each agent is evaluated according to the objective function *f*, and the best-performing agent is identified as

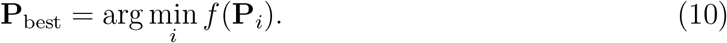

Each agent is also assigned a winding vector **W**_*i*_ = (*w*_*i*,1_, *w*_*i*,2_, …, *w*_*i,d*_), which is initialized to zero. The winding vector records the number of times an agent crosses the boundaries of the search space along each dimension and is used later to refine local search behaviour. Note that, although inspired by the *n*-tuple winding numbers of the torus in topology, which count how many times a path makes a full loop around the space, the winding vector used here is an algorithmic analogue that tracks cumulative boundary crossings along each coordinate direction during the search process. For example, let *γ* be a loop on *T* ^2^ that winds once around each periodic direction of the torus. Then the corresponding homotopy class [*γ*] ∈ *π*_1_ (*T* ^2^) is represented by the winding number pair (1, 1). However, in TSA, if an agent in a rectangular search space crosses the upper boundary once and then the right boundary once, its winding vector is (1, 1) (see Figure 6).

**Figure 6.**
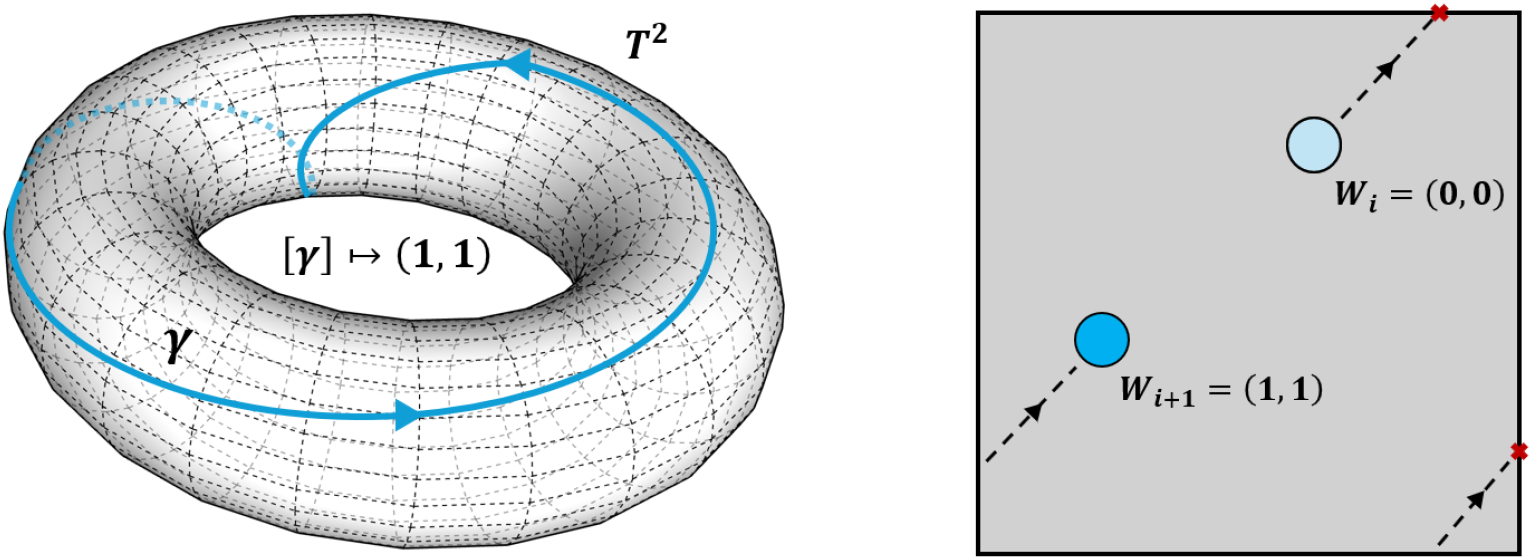
Comparison of the 2-tuple (1, 1) of winding numbers in topology (left) and the winding vector in TSA (right).

### 3.2 Global Search Term

The global search term is responsible for broad exploration of the toroidal search space. At each iteration *t*, the position of **P**_*i*_ is updated according to

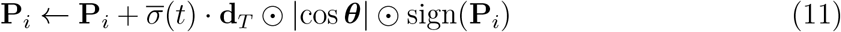

where ***θ*** is a vector of random angles uniformly sampled from [0, *π*], producing values of |cos ***θ***| that are likely to be closer to 1 and thus avoid very small step sizes for exploratory behaviour. The term 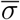 is the weighting factor from the modified sigmoid function (Equation (7)), which activates the global search term in early iterations and suppresses it in late iterations. The toroidal distance **d**_*T*_ is computed using toroidal geometry:

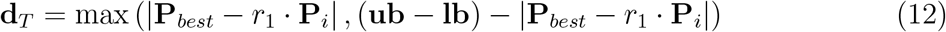

where *r*_1_ ∼ *U* (0, 1) is a random scaling factor. This distance accounts for both the direct distance and the alternate distance which crosses the toroidal boundaries, encouraging agents to explore across the boundaries. Whenever an agent crosses a boundary, it is wrapped back into the domain using the periodic transformation (as in Equation (2))

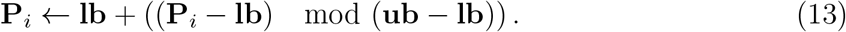

And the corresponding component of the winding vector **W**_*i*_ is updated accorded to

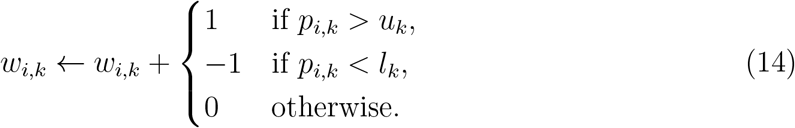

### 3.3 Local Search Term

The local search term takes advantage of the winding vector to refine candidate solutions. Each time an agent crosses a boundary, its winding vector is updated by a signed increment as in Equation (14). Consequently, the *L*^2^-norm of the winding vector represents the net cyclic movement of the agent across all dimensions. Agents with larger winding magnitudes are interpreted as having explored the search space more extensively without locating an optimum. Such agents are therefore encouraged to perform finer local search. For this to occur, larger winding magnitudes lead to smaller step sizes.

The local update term for **P**_*i*_ is therefore given by

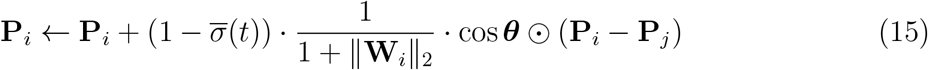

where **P**_*j*_ is a randomly selected agent with *j* =≠ *i*, and ∥**W**_i_∥_2_ is the *L*^2^-norm of the winding vector. The term 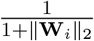 adaptively reduces the step size as the agent traverses boundaries more frequently, promoting more precise local search near promising regions. This local term is weighted by 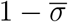, which activates local search in later iterations while deactivating it in early iterations.

### 3.4 Complete Update Mechanism

The complete update rule of TSA combines the global and local search terms. The candidate agent **P**_new_ is therefore generated as follows:

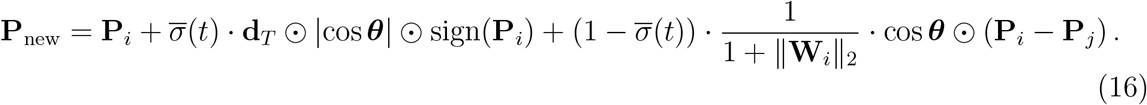

For improved diversity of search, TSA introduces a probabilistic switching strategy. At every iteration *t*, the position of an agent is updated using the toroidal search strategy (Equation (16)) with probability *p*. With probability 1 − *p*, the agent instead follows a direct attraction toward the current best agent:

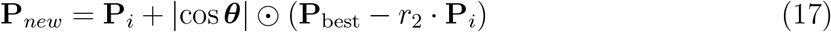

where *r*_2_ ∼ *U* (1, 2) is a random scaling factor. The pseudocode of TSA is summarized in Algorithm 1.

### 3.5 Set-up for Evaluation of Performance

To evaluate the performance of TSA we run two categories of test problems. First, a set of well-known benchmark functions are used to assess the general search capability of the algorithm. These functions include both unimodal and multimodal cases, allowing for comprehensive evaluation of its ability to balance global and local search. Second, TSA is applied to real-world inverse problems in mathematical oncology to demonstrate its practical utility.

To demonstrate performance, we use the benchmark functions from [24, 25] at dimensions *D* = 10, 50, 100, and 500. Fourteen unimodal functions, listed in Table 1, are used to test the performance of TSA at local search and convergence toward the global minimum. Strong performance on these functions suggests that the algorithm is effective at fine-tuning solutions within a single basin of attraction and therefore, that it has strong exploitation capabilities. Similarly, ten multimodal functions, listed in Table 2, are used to test the ability of the algorithm to explore complex landscapes and to escape from local optima. Good performance on multimodal functions suggests strong exploration capabilities, meaning that the algorithm can navigate complex search spaces and avoid premature convergence.

**Table 1:**
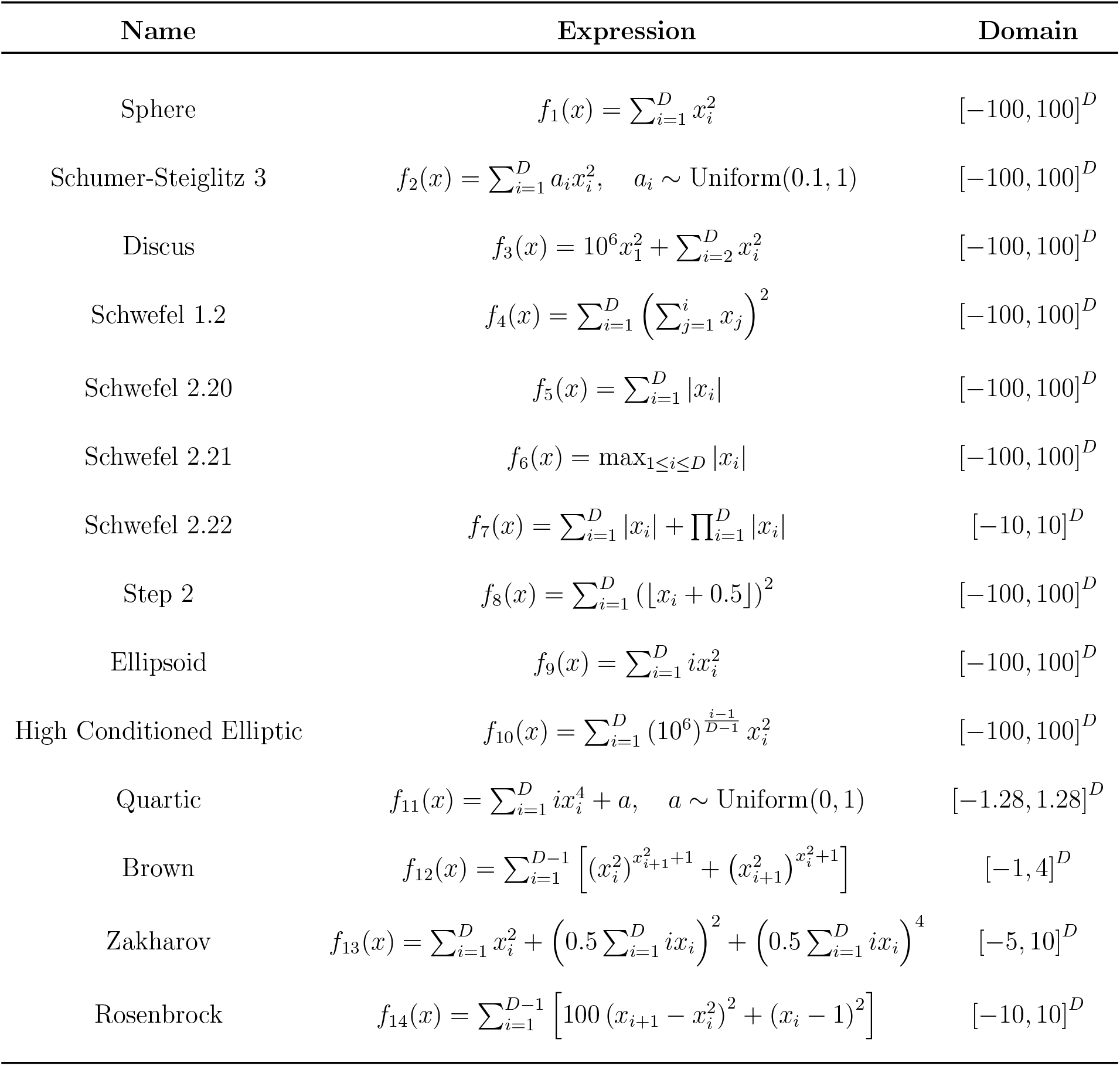
Listing of 14 benchmark unimodal functions. Each function has a minimum of zero, that is, *f*_*i*,min_ = 0 for *i* = 1 … 14.

**Table 2:**
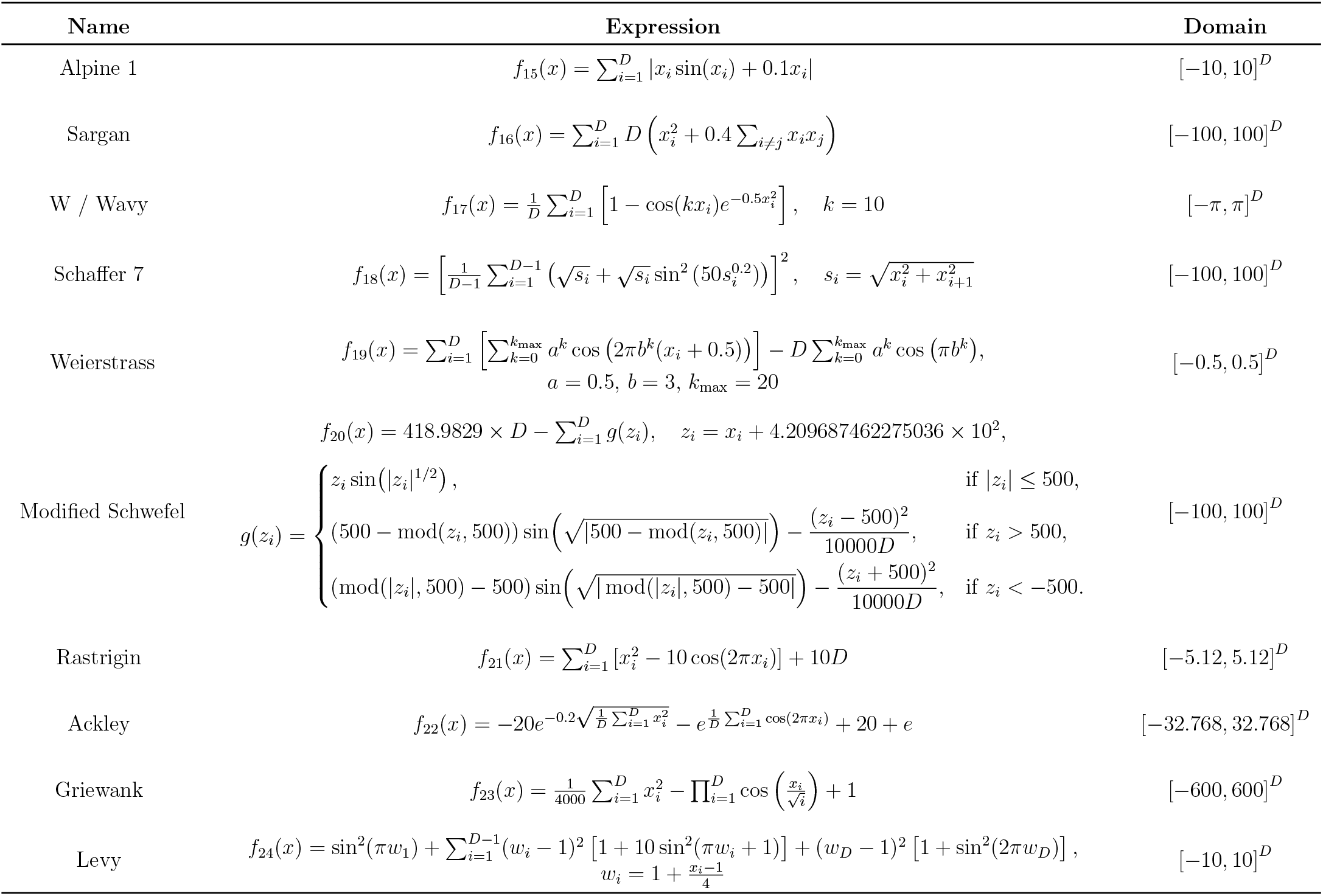
Listing of 10 multimodal benchmark functions. Each function has a minimum of zero, that is, *f*_*i*,min_ = 0 for *i* = 15 … 24.

#### AAlgorithm 1

Pseudocode of Toroidal Search Algorithm (TSA)

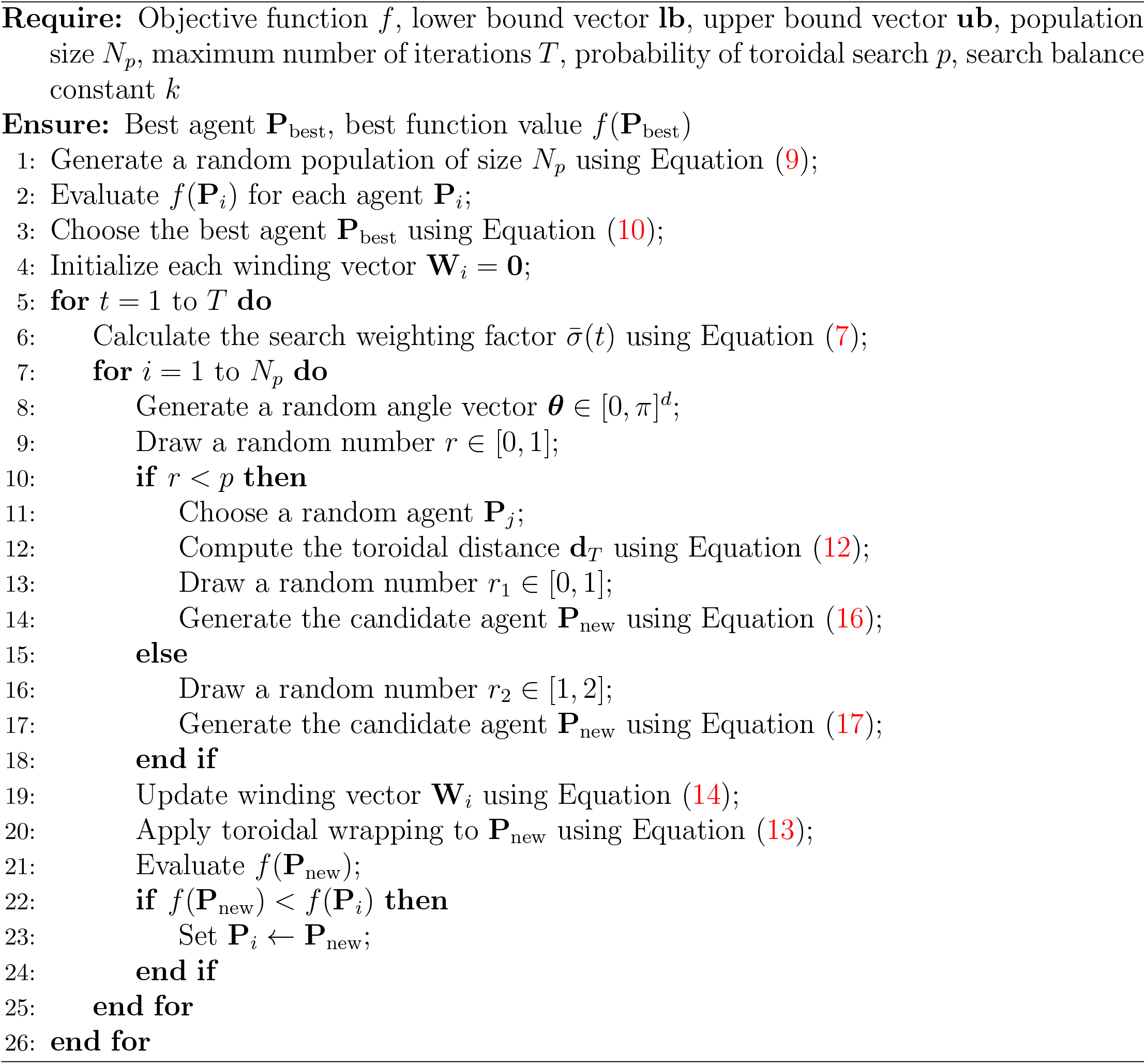

The performance of TSA is compared with five well-established metaheuristic algorithms: PSO, FA, DE, ABC, and TLBO. These algorithms have been widely applied to global optimization tasks and hence are suitable benchmarks for comparison. For each optimization problem, 30 independent runs are conducted to ensure statistical reliability. Each run uses a population size of 30 agents and a maximum of 500 iterations. The reported results include the best, worst, and average function values, the standard deviation, the average running time over all runs, and the convergence curves for each function. Algorithmic parameter values for all algorithms, based on recommendations in the respective literature, are summarized in Table 3.

**Table 3:**
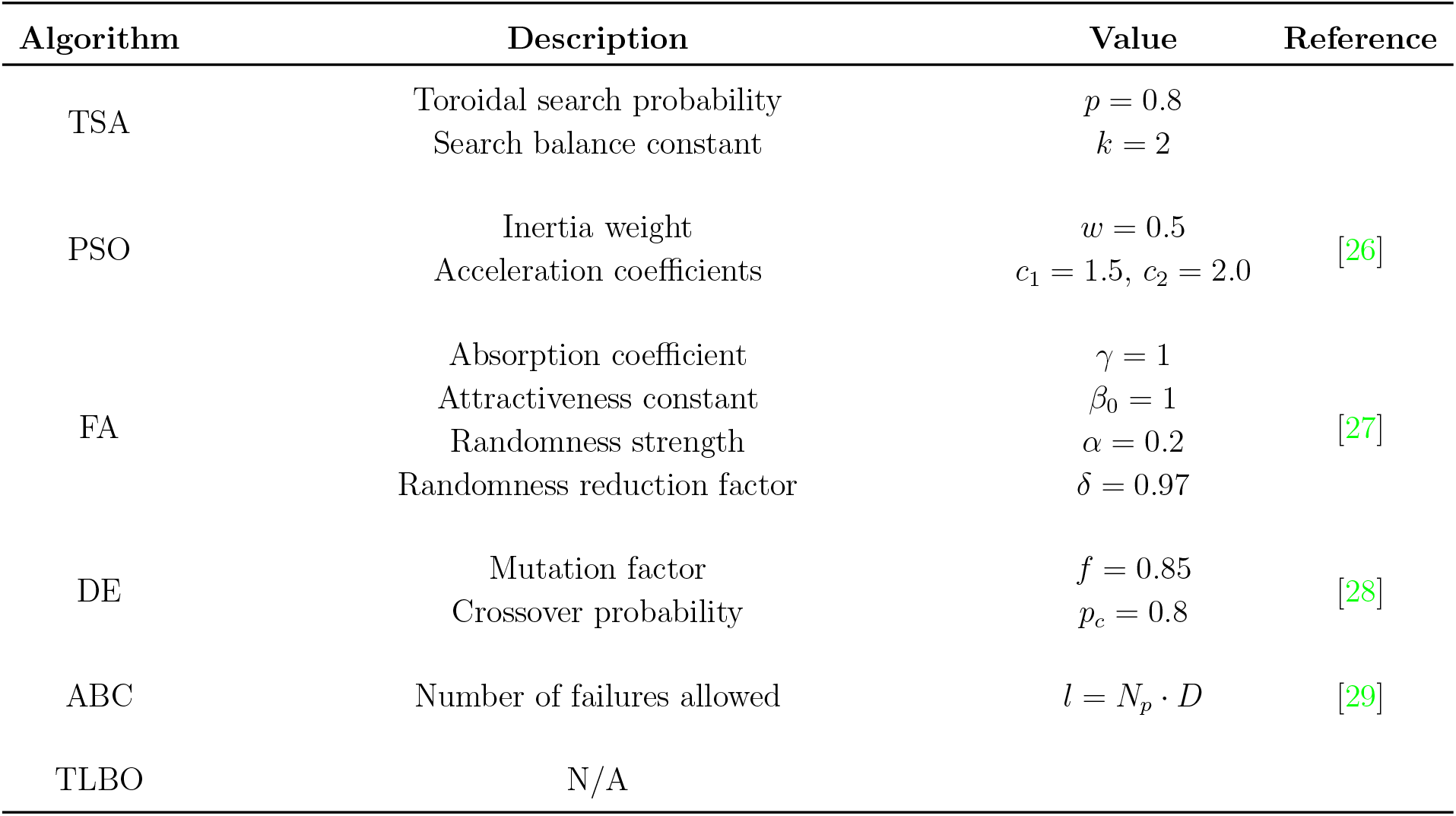
Description of algorithmic parameter values for each algorithm.

| Algorithm | Description | Value | Reference |
| --- | --- | --- | --- |
| TSA | Toroidal search probability | $p = 0.8$ | |
| | Search balance constant | $k = 2$ | |
| PSO | Inertia weight | $w = 0.5$ | [26] |
| | Acceleration coefficients | $c_1 = 1.5, c_2 = 2.0$ | |
| FA | Absorption coefficient | $\gamma = 1$ | [27] |
| | Attractiveness constant | $\beta_0 = 1$ | |
| | Randomness strength | $\alpha = 0.2$ | |
| | Randomness reduction factor | $\delta = 0.97$ | |
| DE | Mutation factor | $f = 0.85$ | [28] |
| | Crossover probability | $p_c = 0.8$ | |
| ABC | Number of failures allowed | $l = N_p \cdot D$ | [29] |
| TLBO | N/A |  |  |

## 4 Results

### 4.1 Optimization of Unimodal Functions

The experimental results in Tables SI.1 – SI.8 demonstrate the strong performance of TSA on 14 unimodal benchmark functions evaluated at dimensions *D* = 10, 50, 100, and 500. Note that when 0.0 is reported in the tables, this indicates that the value has become numerically indistinguishable from zero in double-precision floating-point arithmetic, falling below the smallest positive representable number (5.0 × 10^−324^). TSA attains near-optimal or optimal results on 13 of the 14 functions (with weakest performance on *f*_14_), and consistently outperforms or matches all comparison metaheuristic algorithms (PSO, DE, FA, ABC, and TLBO) on the majority of test functions, particularly as the problem dimensionality increases, see Figure 7. This highlights the robustness and accuracy of TSA in optimizing unimodal landscapes, even in high dimensions.

**Figure 7.**
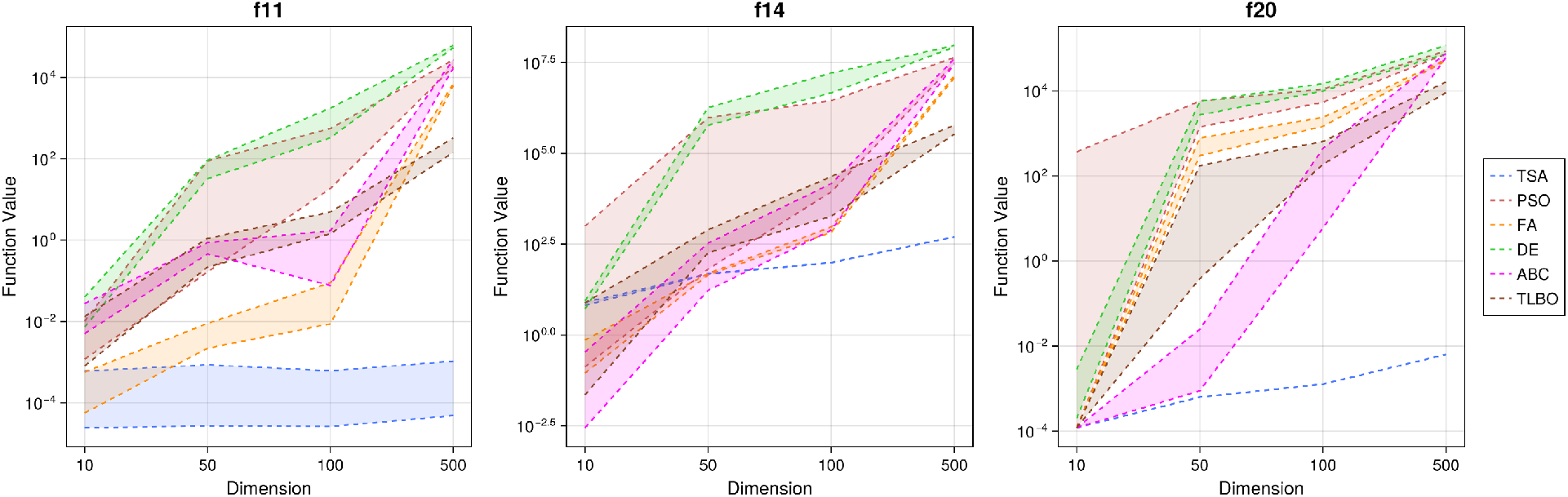
Best–worst ribbon plots of objective function values for *f*_11_, *f*_14_, and *f*_20_ over increasing dimensions. For each algorithm, the dashed curves correspond to the best and worst function values obtained over independent runs, and the shaded region represents the range between them. TSA (blue) exhibits relatively small variation across all runs and a comparatively mild increase in function values as dimension increases, while the other methods show larger spreads and loss of optimality in higher dimensions.

The stability of TSA against stochastic variation is especially evident in functions like *f*_2_ and *f*_11_, where competing algorithms show substantial variability in performance while TSA remains remarkably consistent, see Figure 7. Dimensionality has little effect on the convergence behaviour of TSA as it maintains high accuracy and low variance even in high-dimensional settings. Although minor degradation appears in *f*_11_ and *f*_13_ as dimensionality increases, TSA still surpasses competing methods on these functions. By contrast, competing algorithms experience dramatic degradation in precision with increasing dimensionality.

The most challenging function is *f*_14_, as it is characterized by a narrow, curved valley that makes algorithmic convergence difficult. In low dimensions (*D* = 10), TSA struggles more than other algorithms to efficiently search this landscape. However, as the dimensionality increases (*D* = 50, 100, and 500), the relative performance of TSA improves substantially, while the competing methods suffer severe deterioration, compare for example Tables SI.2 and SI.8, and see Figure 7.

In terms of the average running time per run, TSA is comparable to TLBO in dimensions 10, 50, and 100. It generally requires more time than PSO, DE, and ABC (approximately 1.5 to 2 times longer), but remains significantly faster than FA, often by up to an order of magnitude, while achieving much higher accuracy. Thus, TSA achieves a favourable trade-off between solution quality and computational effort.

The convergence curves for unimodal functions, Figures SI.1 – SI.4, help to illustrate the improved performance of TSA over the other metaheuristics. In all tested dimensions, TSA converges to the global minimum rapidly compared to the competing algorithms. For certain functions, such as *f*_7_, all methods including TSA successfully locate the global optimum in the early stages of the search regardless of dimensionality. For functions like *f*_10_, however, the comparator algorithms converge quickly when *D* = 10 or 50, but take longer at *D* = 100 and 500, due to the added dimensions in the problem. In contrast, TSA continues to converge rapidly in all dimensions for this function, showing robustness to increased dimensionality compared to the other methods. In general, at low dimensions, the comparator algorithms exhibit similar convergence speeds to that of TSA, however, as dimensionality increases, TSA maintains its fast convergence while the others are slower to converge. This highlights the ability of TSA to avoid stagnation and maintain effective search processes even in high-dimensional spaces. Although TSA generally requires a higher computational cost than other algorithms, its early convergence offsets this disadvantage. The algorithm reaches optimal or near-optimal solutions in fewer iterations, which compensates for its greater per-iteration cost.

### 4.2 Optimization of Multimodal Functions

Tables SI.9 – SI.12 summarize the results of TSA on the ten multimodal functions at *D* = 10, 50, 100, and 500, respectively. TSA maintains a clear performance advantage as the dimensionality increases and achieves near-optimal or optimal solutions on eight of the ten functions, demonstrating strong robustness in handling complex search spaces with numerous local minima. While TSA preserves high-quality solutions with only minor degradation at larger scales, the performance of conventional algorithms deteriorates rapidly as the dimension grows.

A particularly notable case is *f*_17_. While TSA produces near-optimal solutions in the low-dimensional case (*D* = 10), it remarkably reaches the exact global minimum in higher dimensions (*D* = 50, 100, 500). Such behaviour is unusual for metaheuristic algorithms as increasing dimensionality typically aggravates the search difficulty due to the curse of dimensionality [30]. The strong performance of TSA in high-dimensional settings can be attributed to its ability to avoid boundary stagnation which intensifies with dimensionality and often destabilizes other methods.

On *f*_24_, TSA performs comparably or slightly worse than the other algorithms (particularly ABC and TLBO) in low dimensions, but performance ranking reverses as dimension increases. While the comparable methods deteriorate sharply in high-dimensional space, TSA maintains stable and competitive performance. This suggests that the robustness and stability of TSA is even more advantageous in high dimensional search spaces.

The benchmark function *f*_20_ presents one of the most deceptive multimodal landscapes, containing numerous local minima that easily trap most metaheuristics. TSA performs exceptionally well on this function, maintaining near-optimal values with all tested dimensions. At *D* = 10, it achieves results close to the theoretical minimum and shows only minor degradation as dimension increases. In contrast, the competing algorithms suffer large performance loss with their errors increasing by several orders of magnitude at high dimensions. The strong performance of TSA can be attributed to the toroidal distance formulation, which allows agents to explore alternate paths across boundaries, effectively escaping local minima and preventing premature convergence.

Regarding computational cost, TSA is generally within a factor of 1.5 to 2 of PSO, DE, and ABC, and comparable to TLBO for dimensions *D* = 10, 50, and 100. It becomes somewhat slower at *D* = 500, but still remains much faster than FA, which is often an order of magnitude slower on the same problem. Given its superior accuracy, this trade-off is reasonable and supports the practicality of TSA for large-scale multimodal optimization.

The convergence behaviour of TSA on multimodal functions, as shown in Figures SI.5 – SI.8, demonstrates its ability to efficiently navigate complex search spaces. Unlike in the unimodal setting where most algorithms exhibit fast convergence in low dimensions, TSA maintains rapid convergence even on multimodal functions at *D* = 10. Across most benchmark functions, TSA either matches or surpasses the convergence speed of other algorithms, reaching high-quality solutions within fewer iterations. A notable exception occurs in *f*_17_ at *D* = 10, where TSA converges more slowly than some comparators. However, the slower convergence is lost at higher dimensions (*D* = 50, 100, and 500) where TSA consistently outperforms the other methods. Again, TSA quickly converges to optimal or near-optimal solutions, while competing methods show delayed or stagnating convergence as dimensionality increases. This pattern highlights the robustness of TSA in high-dimensional, multimodal landscapes. Other algorithms become increasingly susceptible to local optima and search stagnation, but TSA maintains stable and efficient convergence regardless of dimension. Although TSA needs higher computational cost per iteration, its ability to reach optimal solutions early offsets this disadvantage. The fast convergence not only improves solution quality but also compensates for its runtime overhead, making TSA a practical choice for large-scale global optimization.

## 5 Case Study from Mathematical Oncology

### 5.1 Motivation

In mathematical oncology, ordinary differential equation (ODE) models are widely used to describe the temporal dynamics of tumour growth and treatment effects [31]. These models provide a mechanistic framework that connects biological processes such as cell proliferation and drug actions to observable outcomes. Accurate parameterization of such models is essential since the estimated parameters determine the model predictions and thus influence treatment optimization strategies [32].

Parameter estimation is an inverse problem and thus, an optimization problem. The goal is to find the parameter vector 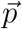 that best fits the model predictions to experimental or clinical data. Consider a tumour growth model of the form

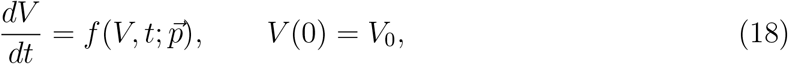

where 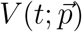 is the tumour volume at time *t* determined by the set of parameters 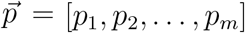. We assume that tumour volumes are measured experimentally at discrete time points, giving a data set {(*t*_1_, *y*_1_), (*t*_2_, *y*_2_), …, (*t*_*n*_, *y*_*n*_)}. A common approach is to estimate 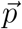 by minimizing the sum of squared errors between the model prediction 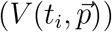 and the observed data (*y*_*i*_):

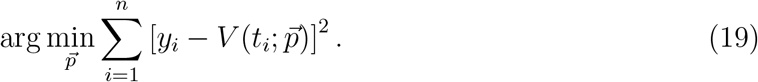

Sometimes, rescaling the problem logarithmically results in better handling of the errors over the full range of data points [33]:

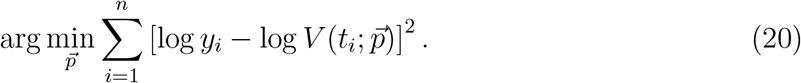

In practice, the objective function, either (19) or (20) or some other defined cost function, typically exhibits a highly complex landscape characterized by strong nonlinearity, nonconvexity, multimodality, and nonseparability [34]. In this study, we use the logarithmic loss function defined in Equation (20), namely the log-transformed least-squares error (LLSE). Even small perturbations in 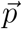 can produce qualitatively different system dynamics, making the identification of a global minimum challenging. Consequently, the choice of optimization method can be significant, as one wishes to balance computational time with adequate global search, finessed the local search, and avoidance of boundary stagnation.

To address these challenges, metaheuristic algorithms have been increasingly adopted for ODE parameterization in cancer modelling [35]. Their success depends largely on their capacity to escape local optima and iteratively refine candidate parameter sets, thus preventing premature convergence. Using the toroidal search mechanism with adaptive search control, we expect TSA to improve convergence reliability and robustness in this setting.

### 5.2 Mathematical Model for Chemotherapy

The effects of chemotherapy on tumour dynamics are commonly modelled by adding a drug-induced cell-kill term to a tumour growth equation. In this case study, we use the log-kill (LK) chemotherapy model, which is widely used and relatively simple. The LK hypothesis assumes that the cytotoxic effect of a chemotherapeutic agent is proportional to tumour size, capturing the idea that a fixed fraction of tumour cells is killed per unit drug exposure [36].

Here, we describe the tumour growth and drug dynamics by the following system of ODEs:

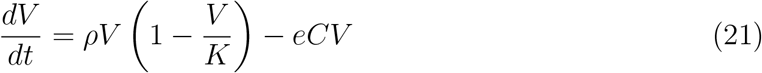

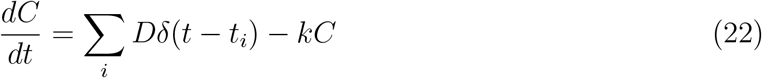

where *V* is the tumour volume, *ρ* is the intrinsic tumour growth rate, *K* is the carrying capacity of the tumour, *C* is the drug concentration from applied dose *D*, the delta function *δ*(*t* − *t*_*i*_) is an instantaneous injection at time *t*_*i*_, *e* is the drug efficacy, and *k* is the drug clearance rate. Tumour growth is described by the logistic growth equation which is a good model for many tumour types [37] that balances simplicity with physiological appropriateness.

### 5.3 Experimental Design

To evaluate the applicability of TSA to ODE parameterization problems in mathematical oncology, we conduct two types of simulation studies: (1) experiments using synthetic tumour–treatment data generated from the LK chemotherapy model described in Equations (21) and (22), and (2) experiments using real-world clinical tumour volume measurements, specifically the prostate cancer dataset from [38], publicly available at https://www.nicholasbruchovsky.com/clinicalResearch.html. The first studies explore the ability of TSA to accurately reconstruct underlying biological parameters from noisy and sparsely sampled data, as well as to investigate the influence of population size on performance. The second studies assess the ability of TSA to find physiologically plausible parameters from clinical data and thereby generate meaningful predictive tumour trajectories.

#### 5.3.1 Synthetic Data Experimental Setup

We first generate noise-free tumour volume trajectories by numerically solving the ODE system for the LK chemotherapy model using the true parameter values listed in Table 4. Tumour dynamics are simulated over the time window *t* ∈ [0, 50].

**Table 4:** True model parameter values used in the synthetic dataset for the LK chemotherapy model, Equations (21) and (22).

| Parameter | Description | Value | Unit |
| --- | --- | --- | --- |
| $\rho$ | Tumour growth rate | 0.1 | day<sup>-1</sup> |
| $K$ | Carrying capacity of tumour | 2000 | mm<sup>3</sup> |
| $e$ | Drug efficacy | 0.06 | kg/(mg·day) |
| $k$ | Drug clearance rate | 0.5 | day<sup>-1</sup> |
| $d$ | Treatment dose | 10 | mg/kg |
| $t_i$ | Delivery schedule of treatment | 5, 15, 35 | day |
| $V(0)$ | Initial tumour volume | 100 | mm<sup>3</sup> |
| $C(0)$ | Initial drug concentration | 0 | mg/kg |

To mimic measurement variability typically observed in clinical data, we add 5% Gaussian noise to each datapoint. The observed synthetic measurements *y*_*i*_ are thus defined by

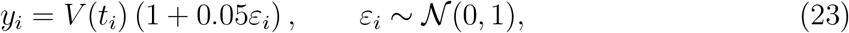

where *ε*_*i*_ are independent standard normal random variables. Twenty-five observation times are then selected uniformly at random from the full simulated trajectory to approximate sparsely sampled clinical data. The synthetic dataset is shown in Figure 8.

**Figure 8.**
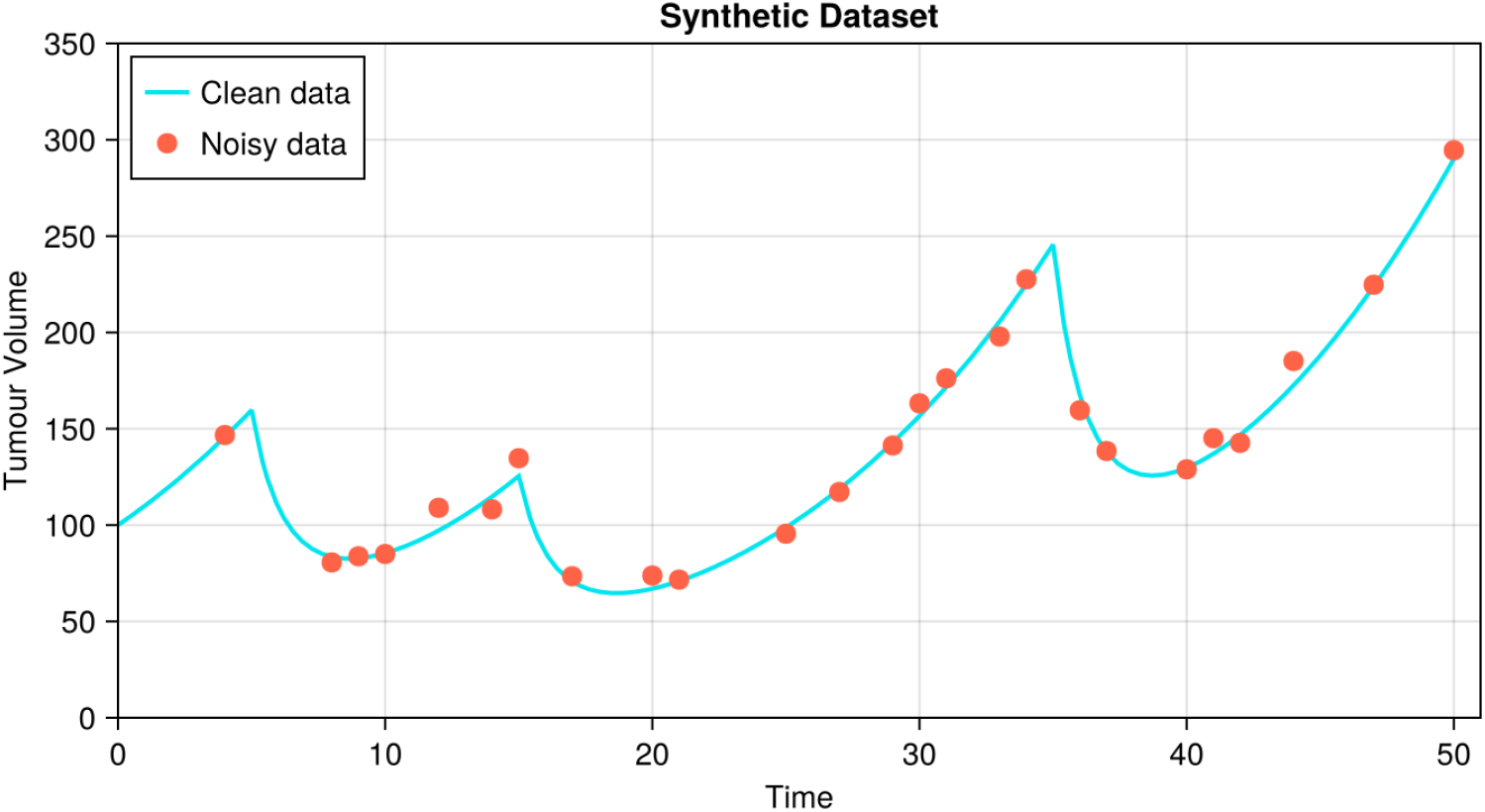
Synthetic tumour volume data generated from the LK chemotherapy model. Clean (noise-free) model trajectory (blue curve), and the corresponding sparsely-sampled and noisy observations used for parameter estimation (red circles).

TSA is used to estimate the following four parameters by fitting the model simulation to the synthetic data:

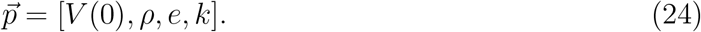

The remaining parameters are fixed due to experimental design choices. Carrying capacity is fixed at *K* = 2000 since the parameter is known to be practically non-identifiable when tumour datasets are sparse or contain only early-stage measurements, as reported in [33, 39]. Parameter estimation is carried out by minimizing the LLSE. Lastly, the search domain for the estimated parameters is defined as

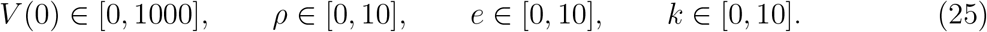

To assess the impact of population size on optimization performance, TSA is tested with *N*_*p*_ = 5, 10, 15, and 20. Each configuration is run 30 times with each run consisting of 5000 iterations to ensure enough search depth to observe the effect of population size.

Moreover, to examine the robustness of TSA to increasing data uncertainty, we regenerate synthetic datasets with Gaussian noise levels of 5%, 10%, 15%, and 20%, and repeat the parameter estimation for each noise level using TSA and PSO with *N*_*p*_ = 5. To quantify how estimation accuracy degrades with noise, we compute the average relative percentage error (ARPE) of the estimated parameters:

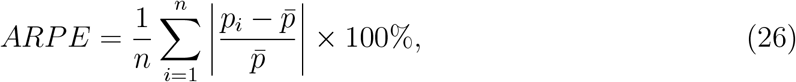

where *n* is the number of runs (i.e., *n* = 30), *p*_*i*_ is the estimated parameter value from the *i*-th run, and 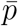 is the corresponding true parameter value.

#### 5.3.2 Clinical Dataset Experimental Setup

The prostate cancer clinical dataset, recently analyzed in [40], contains patient treatment observations such as (1) patient number, (2) observation date, (3) Cyclophosphamide (CPA) dose, (4) Leuprolide (LEU) dose, (5) prostate-specific antigen (PSA) level, (6) testosterone level, (7) cycle number, (8) treatment status, (9) day number, and (10) an alternative day number. Here, we consider three reported variables: PSA level, CPA dose, and the alternative day number. PSA is a serum biomarker routinely used in prostate cancer management and is used here as a biomarker for tumour burden. This assumption is consistent with a previous modelling study [40], which used PSA dynamics as a surrogate for tumour growth and treatment response. To allow comparison between patients, PSA levels are normalized to the baseline measurement at the start of therapy. CPA is an alkylating chemotherapeutic drug widely used in prostate cancer therapy.

For demonstration purposes, we select three patients (numbers 39, 62, and 79) from the dataset as their PSA trajectories exhibit three different tumour response patterns to the CPA treatment (see Figure 9). For each patient, we estimate the LK chemotherapy model parameters (*ρ, e, k*) within the following search domains:

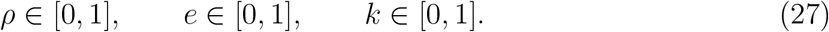

**Figure 9.**
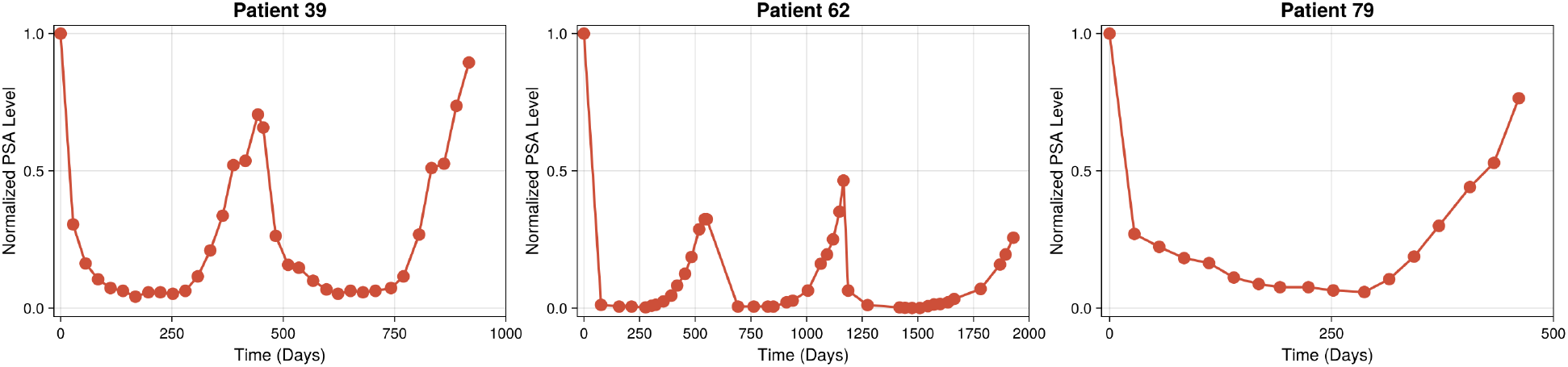
Normalized PSA trajectories for three selected patients (39, 62, and 79) from the Bruchovsky prostate cancer clinical dataset.

Because PSA values are normalized here, the initial PSA level is fixed at *V* (0) = 1. The carrying capacity is fixed at *K* = 1.5.

To test TSA, we examine the question of how much patient-specific data is needed to obtain parameter estimates that yield accurate future predictions. To do this, we first fix the number of data points used for fitting, estimate the parameters, and then predict the remaining future PSA levels given the treatment plan. Estimation error is measured using the mean squared error (MSE):

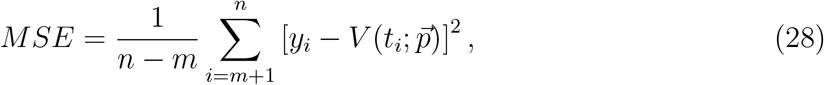

where *m* denotes the number of data points used for parameter estimation. We increase *m* to evaluate how prediction quality improves as more information becomes available and to determine the amount of data each algorithm requires for reliable parameterization.

For all experiments in this section, we use *N*_*p*_ = 20, 500 iterations, and 10 independent runs. Reported parameter value estimates are the average over the 10 runs. An estimate is classified as “good” if the corresponding MSE is less than or equal to 0.1, which ensures that the resulting trajectories closely align with the data.

### 5.4 Results

#### 5.4.1 Parameter Estimation using Synthetic Data

We use TSA and the comparator algorithms to estimate the parameters of the LK chemotherapy model from the synthetic data at different population sizes. Table SI.13 lists the LLSEs, and Table SI.14 lists the estimated parameter values, for each algorithm and population size. The results show that TSA consistently outperforms the competing algorithms over all tested population sizes. A notable finding is that TSA achieves essentially identical best, worst, and mean errors for every value of *N*_*p*_, indicating that its performance is remarkably robust to the number of agents. Even with *N*_*p*_ = 5, TSA reliably converges to the optimal solution, whereas the other algorithms require substantially larger populations (or more iterations) to reach comparable accuracy. For instance, TLBO succeeds reliably with *N*_*p*_ ≥ 15, and the other algorithms require even larger populations. The good results of TSA can be attributed to its inherent strength in high-dimensional search spaces. Reducing the number of agents effectively increases the relative dimensionality of the problem, making the search more challenging. Nevertheless, TSA maintains strong performance due to its toroidal exploration mechanism and robustness against dimensionality-induced degradation. In contrast, the other algorithms exhibit substantial dependence on population size with larger *N*_*p*_ generally improving their best performance but still suffering from large variability and frequent failures, as shown in Figure 10.

**Figure 10.**
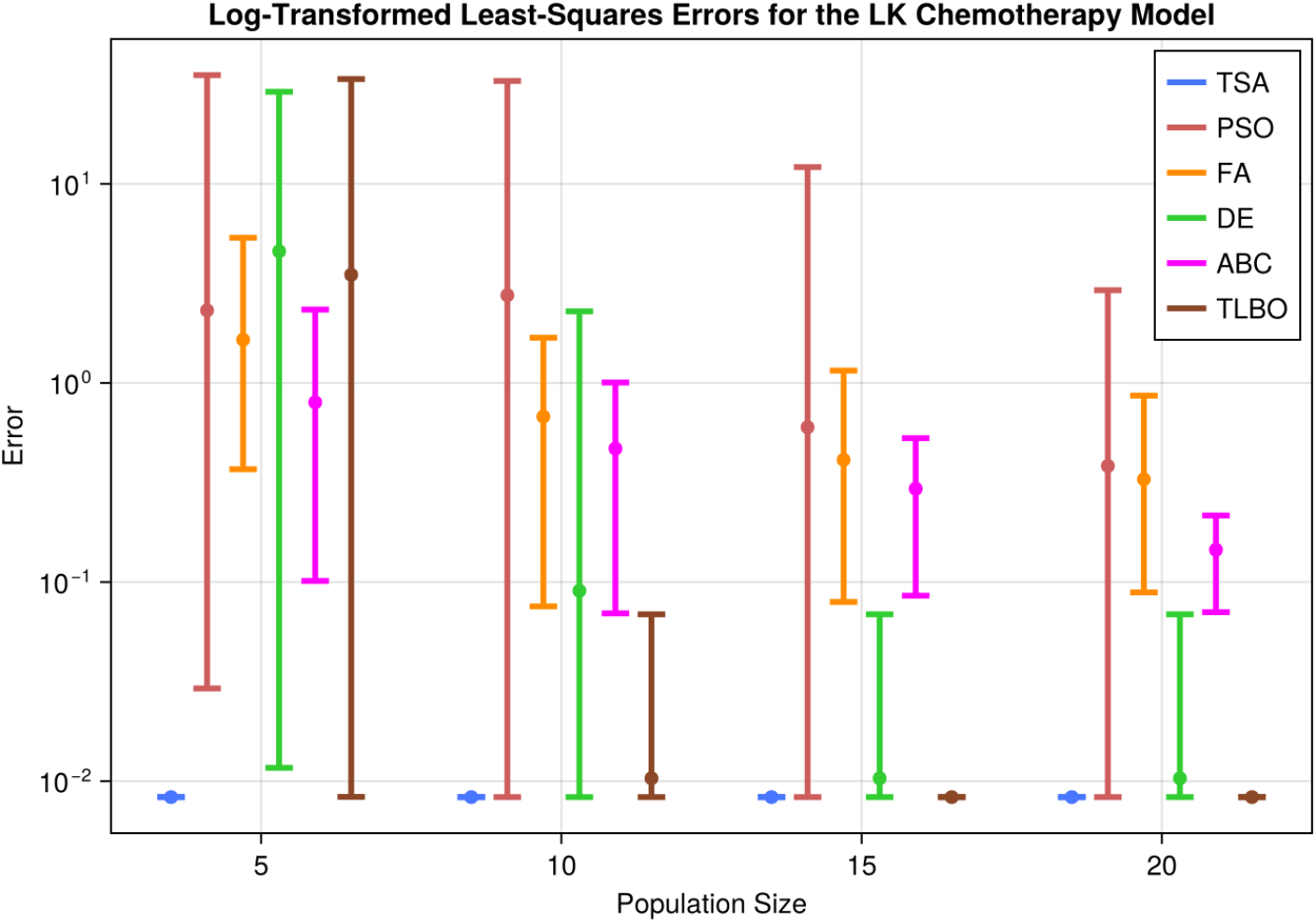
Best and worst values of the LLSEs (dots indicate the mean) from LK chemotherapy model parameter estimation using the synthetic data for population sizes *N*_*p*_ = 5, 10, 15, and 20 for all algorithms.

A distinguishing characteristic of TSA is its extremely low variance across the runs. The standard deviations of TSA fitting errors are several orders of magnitude smaller than those of the competing methods, reflecting a tendency of these methods to get stuck in poor regions of the parameter space. These failures can be significant to ODE parameterization and predictions as these ill-fitting values may cause numerical stiffness or unstable trajectories, increasing computational cost, and degrading reliability. TSA avoids this issue entirely, as even its worst-performing run recovers parameter values that closely match the true values and do not collapse toward boundary values or unrealistic parameter combinations.

To further illustrate the exceptional stability of TSA, we simulate the LK model using all 30 sets of parameter estimates obtained with *N*_*p*_ = 5. As shown in Figure 11, the resulting tumour–volume trajectories overlap almost perfectly, producing what appears to be a single curve despite being the overlap of 30 different simulations. TSA is remarkably stable and robust even when using very small population sizes.

**Figure 11.**
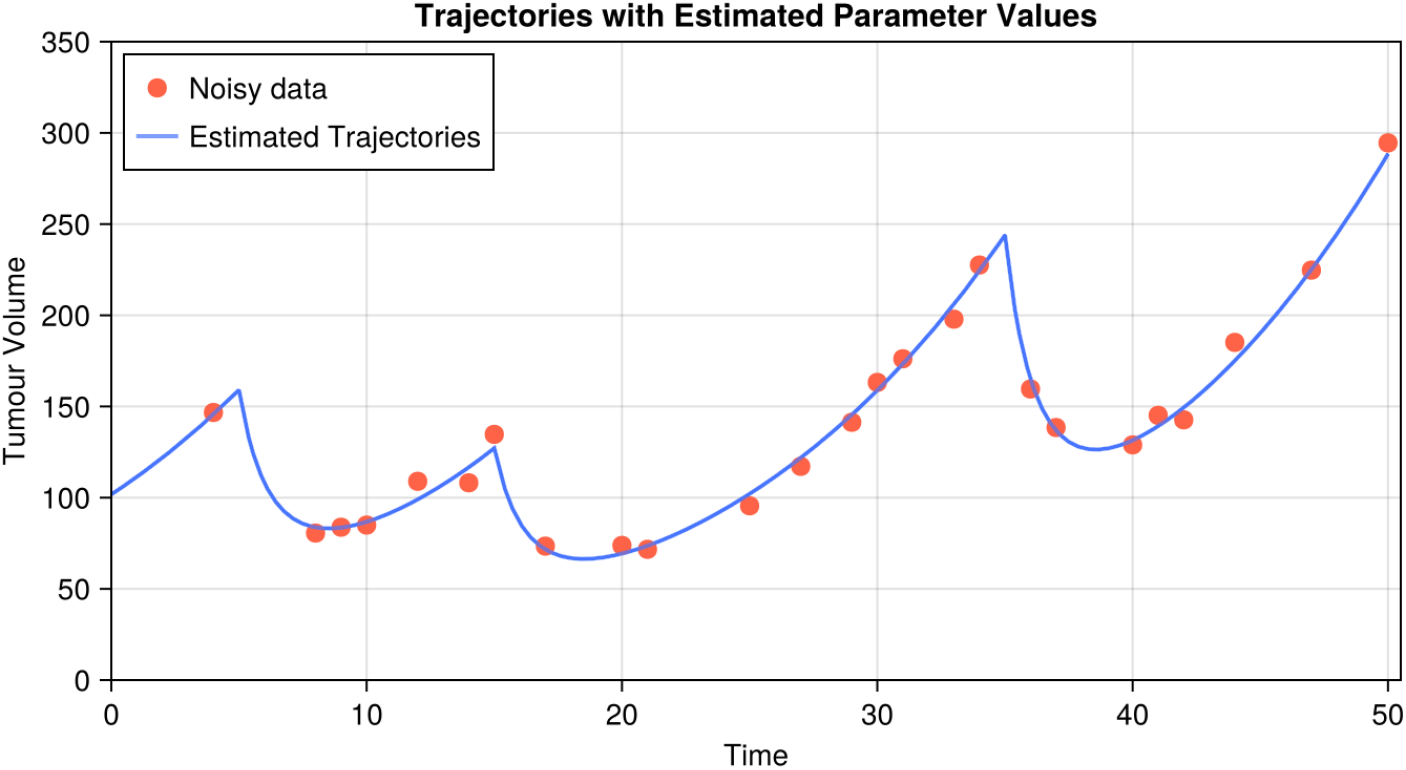
Realizations of the LK model using the 30 parameter sets estimated by TSA with *N*_*p*_ = 5.

Surprisingly, TSA achieves the lowest or comparable computation times relative to all the tested algorithms on this ODE parameter estimation problem (see Table SI.13), even though it generally has a higher per-iteration computational cost in the benchmark tests (see Tables SI.5–SI.8 and SI.9–SI.12). In these tests, TSA typically requires approximately 1.5–2 times the running time of DE, while remaining comparable to TLBO in most cases. On the other hand, in the ODE parameter estimation problem, TSA achieves the lowest overall runtime in most cases and remains comparable to DE in the remaining cases.

This apparent inversion arises from the nature of ODE-based parameter estimation problems, where algorithms that explore poor parameter regions can force the ODE solver into stiff or dynamically unstable regions, increasing computation time. TSA, however, converges rapidly toward well-behaved parameter regions, as illustrated in Figure 12, and thereby avoids such instability. Early convergence in TSA significantly reduces the number of expensive ODE evaluations, giving it a practical speed advantage over the other comparators. Moreover, although DE is also among the fastest algorithms in this experiment, it often converges to suboptimal solutions (see Figure 10). This behaviour implies that it is generally effective at global search and can reliably locate the region containing the optimal solution, but it lacks strong local search capabilities and therefore struggles to refine solutions once it reaches that region. In contrast, TSA maintains both robust global search and precise local search, enabling it not only to find the correct region but also to accurately converge to the optimal parameter values.

**Figure 12.**
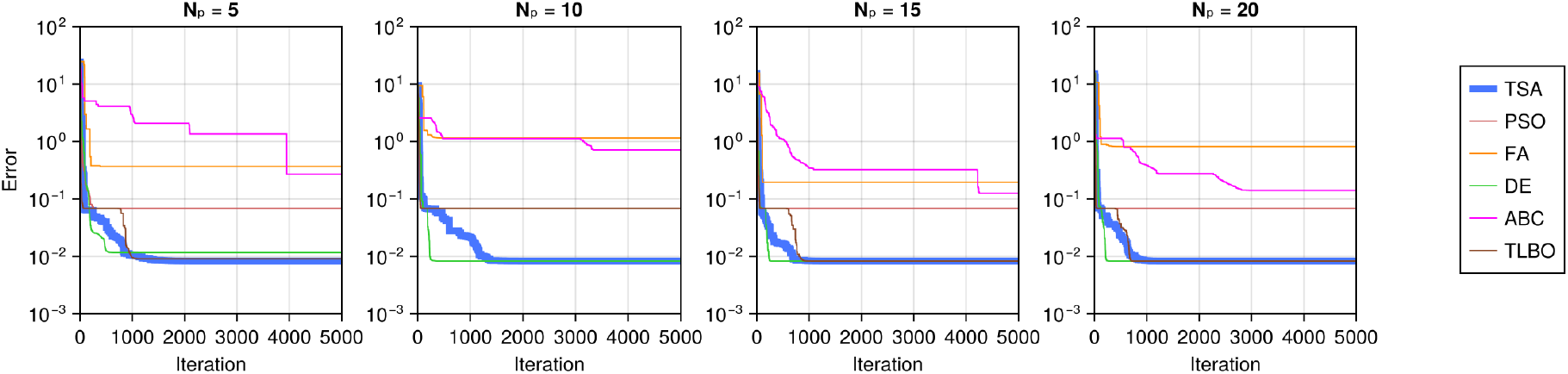
Convergence behaviour of TSA and the competing algorithms for different population sizes (*N*_*p*_ = 5, 10, 15, and 20).

Figure 13 summarizes the average relative percent error (ARPE, Equation (26)) of the estimated parameters for TSA and PSO under increasing noise conditions. As expected, ARPE increases monotonically with noise for all TSA parameter estimates, indicating that accuracy deteriorates as data uncertainty grows. Nevertheless, the overall error levels for TSA remain moderate even at 20% noise. Parameter *k* shows the greatest ARPE and is therefore more sensitive to noise than the other parameters, such as *V* (0), which is relatively unaffected. On the other hand, PSO shows ARPE values that are several orders of magnitude larger than those of TSA. The ARPE curves for PSO are high (around 10^3^) and approximately flat, not because the algorithm is robust to noise, but because its estimation error is already extremely large even at 5% noise. It is important to note that the performance of PSO is strongly influenced by the small population size used here (*N*_*p*_ = 5).

**Figure 13.**
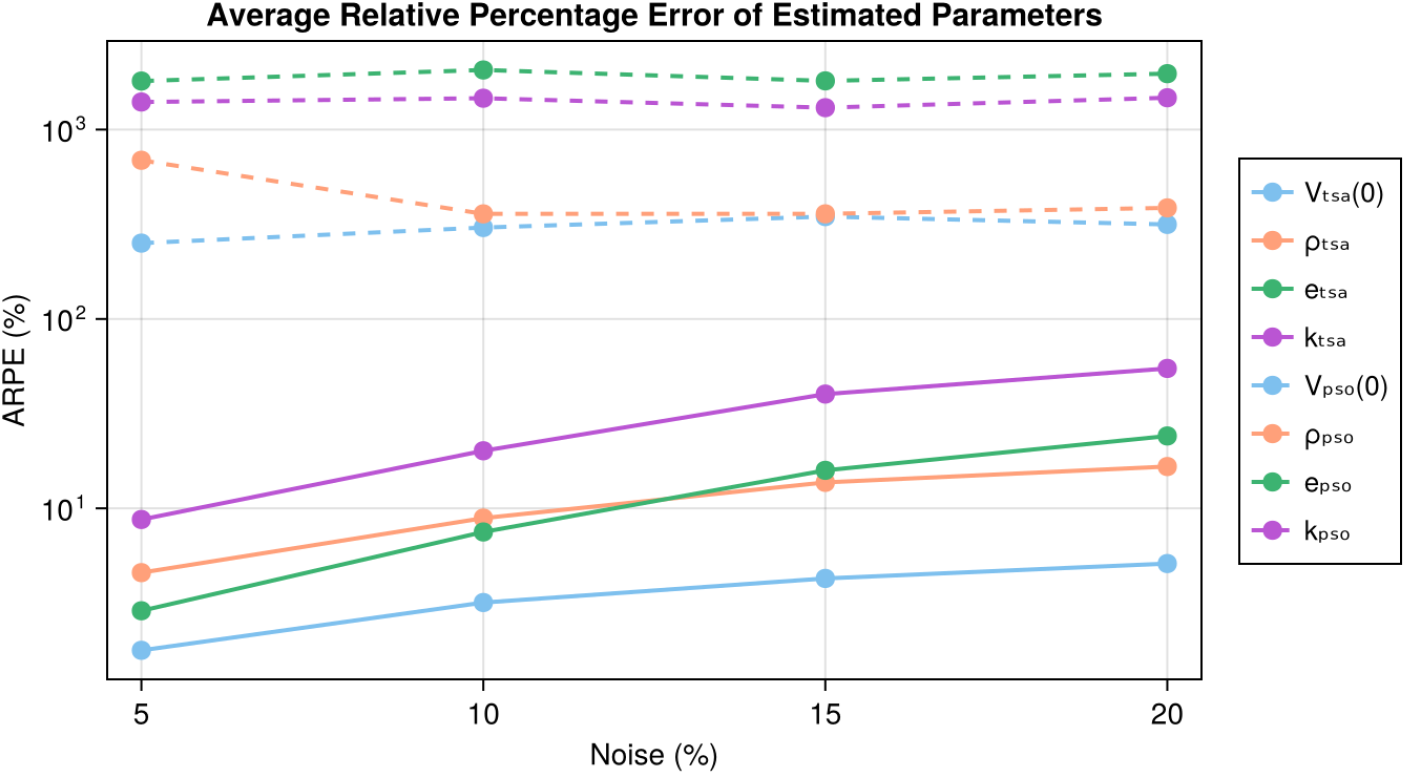
Average relative percentage error of the estimated parameters for TSA and PSO with *N*_*p*_ = 5 under increasing noise levels (5%, 10%, 15%, and 20%). TSA estimates solid lines and PSO estimates dashed lines.

#### 5.4.2 Parameter Estimation using Clinical Data

Table 5 summarizes the prediction performance of the tested algorithms for three prostate cancer patients when increasing numbers of data points, *m*, are used for parameter estimation. The reported MSEs are computed over the remaining data points not used in the fitting. The results demonstrate that prediction accuracy depends not only on the quantity of available data, but also on the complexity of the patient-specific tumour dynamics, and on the robustness of the optimization algorithm.

**Table 5:** MSEs from LK chemotherapy model predictions for Patients 39, 62, and 79 obtained by the tested algorithms with increasing numbers of fitted data points (*m*).

| Patient | $m$ | TSA | PSO | FA | DE | ABC | TLBO |
| --- | --- | --- | --- | --- | --- | --- | --- |
| 39 | 5 | 0.12032 | 1.45568 | 0.37299 | 0.12005 | 1.46865 | 1.58214 |
|  | 10 | 0.02466 | 0.44533 | 0.45031 | 0.13193 | 0.49462 | 0.28070 |
|  | 15 | 0.05481 | 0.87833 | 0.45682 | 0.06667 | 0.43222 | 1.47543 |
|  | 20 | 0.00563 | 0.45451 | 0.45252 | 0.00612 | 0.35493 | 1.59109 |
|  | 25 | 0.00769 | 0.49037 | 0.54170 | 0.00787 | 1.14847 | 1.38478 |
| 62 | 5 | 1.09642 | 1.90834 | 1.90926 | 1.90746 | 1.25466 | 1.90862 |
|  | 10 | 0.56077 | 1.87109 | 1.87206 | 1.86995 | 1.28649 | 1.87131 |
|  | 15 | 0.38072 | 1.92646 | 1.92799 | 1.92549 | 1.13898 | 1.92710 |
|  | 20 | 0.13794 | 1.86103 | 1.86103 | 1.86103 | 1.27278 | 1.86104 |
|  | 25 | 0.09097 | 1.86243 | 1.86243 | 1.86243 | 1.22945 | 1.86243 |
|  | 30 | 0.09719 | 2.06742 | 2.06742 | 2.06742 | 0.57845 | 2.06742 |
|  | 35 | 0.09483 | 1.94707 | 1.94707 | 1.94707 | 1.10026 | 1.94707 |
|  | 40 | 0.09319 | 1.62325 | 1.62325 | 1.62325 | 1.62218 | 1.62325 |
| 79 | 5 | 0.31497 | 1.65075 | 0.64540 | 1.48793 | 1.58861 | 1.65125 |
|  | 10 | 0.00672 | 1.39787 | 1.10458 | 0.00658 | 1.39787 | 0.57141 |
|  | 15 | 0.00683 | 0.74135 | 0.74135 | 0.00675 | 0.74135 | 0.74135 |

As shown in Figure 9, the PSA trajectories of the three patients exhibit different dynamics. Patient 79 has a single valley, Patient 39 two valleys, and Patient 62 three valleys. These valleys correspond to cycles of tumour growth and treatment-induced regression, which increases the complexity of the parameter-estimation problem.

For Patients 39 and 79, only TSA and DE are able to produce good estimations according to the criterion MSE ≤ 0.1 (see Figure 14). For Patient 39, which has two cycles, TSA achieves the desired accuracy when *m* = 10, while DE requires at least *m* = 15. For Patient 79, with one cycle, both algorithms require at least *m* = 10, but TSA produces a substantially lower MSE than DE when *m* = 5. In contrast, PSO, FA, ABC, and TLBO fail to provide reliable predictions for either patient. Although these algorithms occasionally find a good estimate among the 10 runs, they predominantly suffer from premature convergence and fail to escape poor regions of parameter space. Since the population size (*N*_*p*_ = 20) and the number of iterations (500) are sufficient to estimate the three model parameters, as demonstrated by TSA and DE, these failures cannot be attributed to an insufficient number of agents or iterations, but rather to a lack of robustness within the algorithms to navigate the complex landscapes.

**Figure 14.**
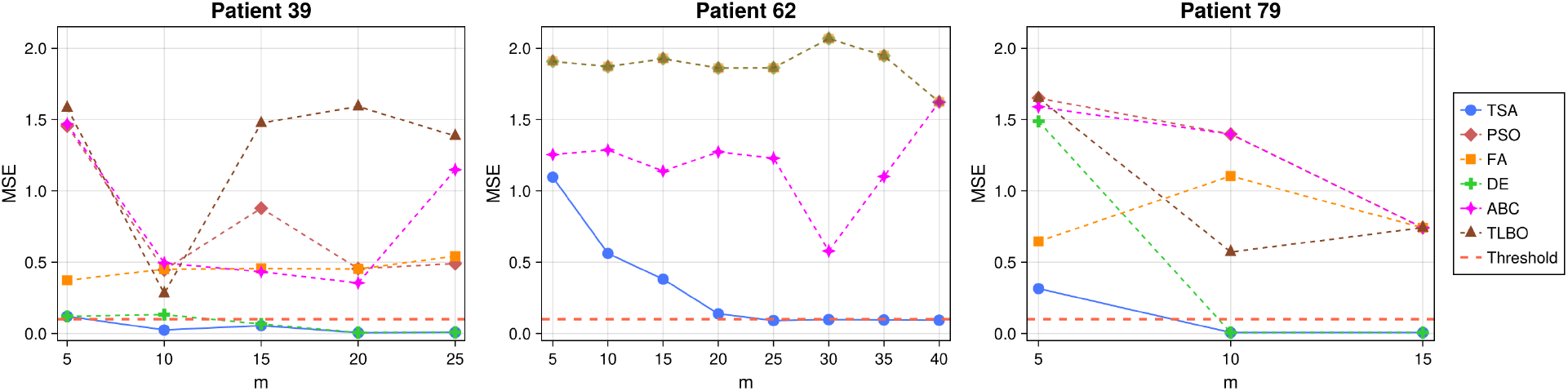
MSE curves of LK chemotherapy model predictions for Patients 39, 62, and 79 as the number of data points used in the estimation, *m*, increases.

For Patient 62, which has three cycles, TSA is the only algorithm capable of producing good estimates. Even DE, which performed competitively for the other tested patients, fails to achieve acceptable accuracy regardless of the number of data points used. On the other hand, TSA successfully achieves MSE ≤ 0.1 once *m* ≥ 25. The increased data requirement reflects the inherent difficulty of the problem rather than a limitation of TSA, as additional information is necessary to reliably estimate model parameters in the presence of multiple treatment–response cycles.

These results further indicate that increasing the number of fitting data points does not necessarily lead to improved prediction accuracy unless the optimization algorithm is sufficiently robust. As illustrated in Figure 14, TSA and DE benefit from additional data, exhibiting decreasing MSE as *m* increases. In contrast, the other algorithms often show little improvement with the use of additional data, suggesting that for algorithm that lack robustness and have difficulty converging, increasing the amount of data does not prevent premature convergence or poor exploration of the parameter space. Consequently, the number of available data points becomes a meaningful factor only when the underlying optimization method is stable.

Table 6 lists the estimated LK model parameters obtained using all tested algorithms. The value *m* = 10 was used for Patients 39 and 79 and *m* = 25 for Patient 62. These *m* values correspond to the smallest number of data points needed for TSA to achieve our prediction accuracy (MSE ≤ 0.1). Although TSA yields parameter estimates that result in model predictions that reasonably agree with the data, the competing algorithms yield parameter estimates that predict dynamics with considerable variability and unphysiological responses, reflecting sensitivity to premature convergence and insufficient exploration in these algorithms (see Figure 15).

**Table 6:**
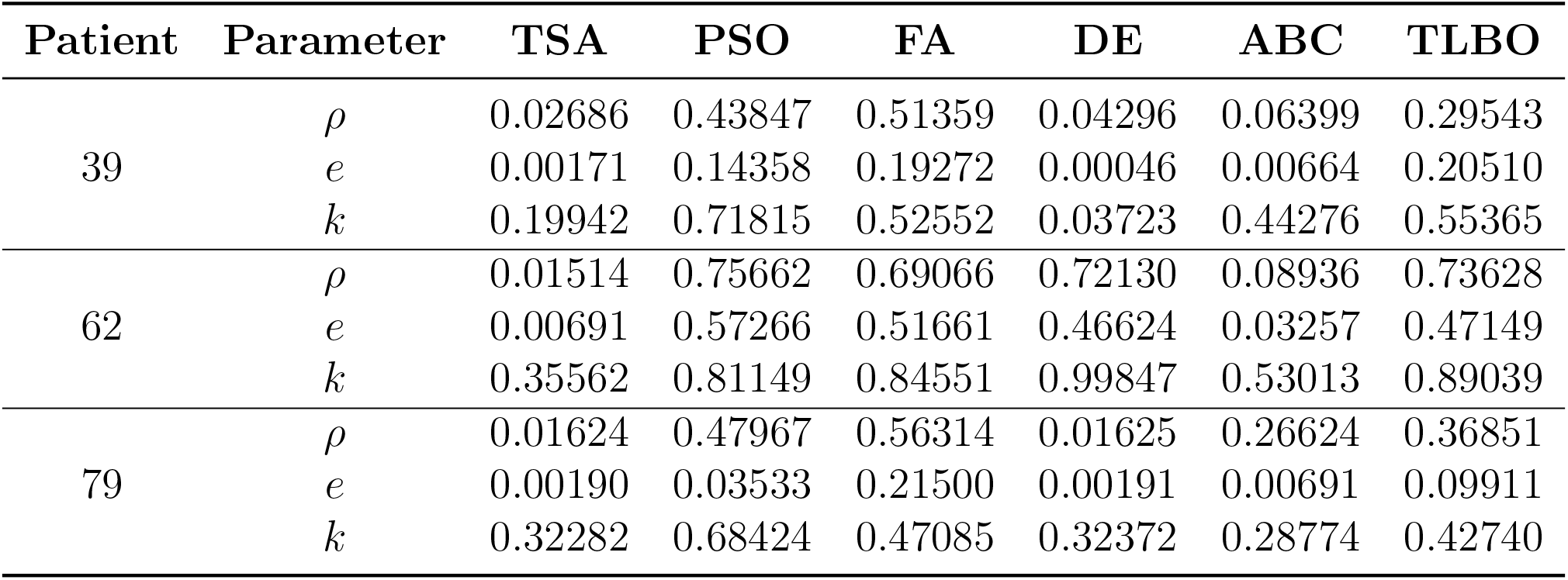
Estimated LK chemotherapy model parameters (*ρ, e, k*) for Patients 39, 62, and 79 obtained using the tested algorithms. Estimates use *m* = 10 data points for Patients 39 (two cycles) and 79 (one cycle) and *m* = 25 for Patient 62 (three cycles).

**Figure 15.**
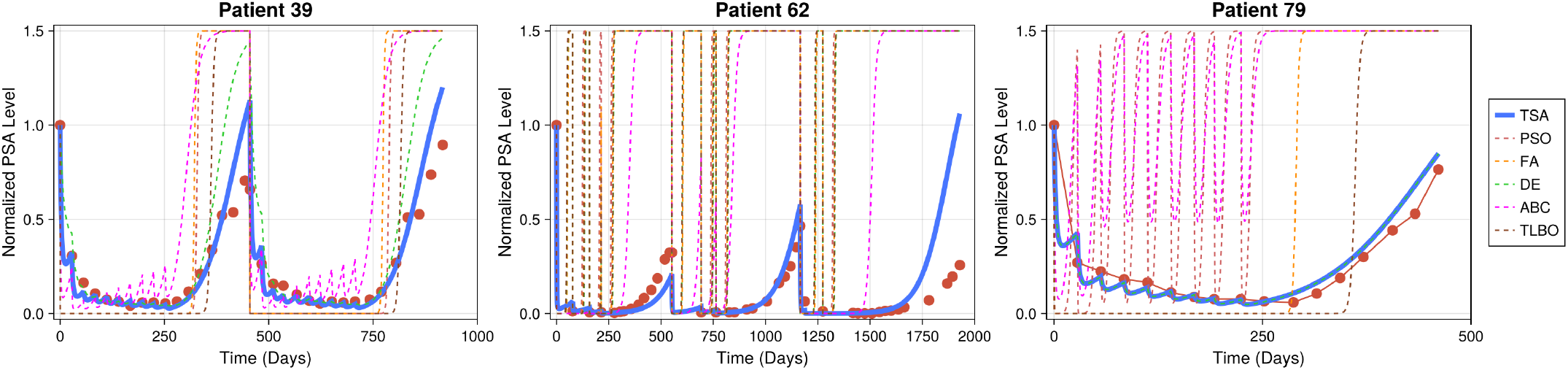
Simulations of the LK chemotherapy model for Patients 39, 62, and 79 using the parameter estimates reported in Table 6.

An important practical insight from these results is the role of algorithmic stability. For metaheuristic optimization, there is no universal rule for selecting the population size or the number of iterations required to avoid premature convergence [41, 42]. This uncertainty is especially critical in ODE-based parameter estimation where each evaluation can be expensive. Therefore, algorithms that exhibit high variability over the individual runs, require extensive tuning and repeated executions, substantially increasing both computational cost and user effort. TSA demonstrates consistently good stability, producing low-variance results across all patients and number of data points. This reliability allows TSA to generate accurate and physiologically plausible parameter estimates with minimal trial-and-error adjustment, making it particularly well suited for ODE parameterization problems.

## 6 Conclusion

We present TSA, a topology-inspired metaheuristic aimed at reducing boundary-related stagnation and improving scalability in high-dimensional optimization. By treating the search space as a torus, TSA removes artificial edges and allows agents to move smoothly and continuously across boundaries. Unlike conventional wrapping strategies applied only after boundary violations occur, this toroidal structure is built directly into the search process.

A key feature of TSA is the use of the toroidal distance to guide global search, which naturally generates winding numbers that track how extensively agents traverse the toroidal search space. These winding numbers are then used to regulate local search by adaptively scaling movement, so agents that have already explored widely, take progressively smaller steps and shift toward local refinement. Together with a modified sigmoid control that governs the timing of global and local search, this mechanism allows TSA to transition from broad exploration to focused exploitation while maintaining a stable balance between search diversity and convergence.

Extensive tests on a wide range of unimodal and multimodal benchmark functions show that TSA typically outperforms, and in some cases, matches, the well-established metaheuristic algorithms we tested against. Its performance remains stable as problem dimensionality increases, with only minor degradation even in very high-dimensional cases. In addition, TSA exhibits low run-to-run variability, reflecting stable convergence behaviour.

We applied TSA to the parameterization problem for an ODE-based tumour growth and chemotherapy model in mathematical oncology. In both synthetic and clinical datasets, TSA reliably identified physiologically plausible parameter sets with substantially lower variance than competing methods. The algorithm proved particularly effective in challenging scenarios involving noisy data and complex tumour–treatment dynamics, where many traditional metaheuristics exhibited premature convergence or instability. These results highlight the strength and practical applicability of TSA in scientific domains where robustness and stability is critical.

## Data Availability

Code for TSA is available at https://github.com/ChanginOh/TSA. The clinical dataset is available at https://www.nicholasbruchovsky.com/clinicalResearch.html.

## Author Contributions

Conceptualization (CO, KPW), Formal Analysis (CO), Funding Acquisition (KPW), Investigation (CO, KPW), Methodology (CO, KPW), Project Administration (CO, KPW), Software (CO), Supervision (KPW), Validation (CO, KPW), Visualization (CO), Writing-Original Draft (CO), Writing-Review and Editing (CO, KPW).

## Supplemental Information

### SI.1 Results for Unimodel Benchmark Functions

**Table SI.1:**
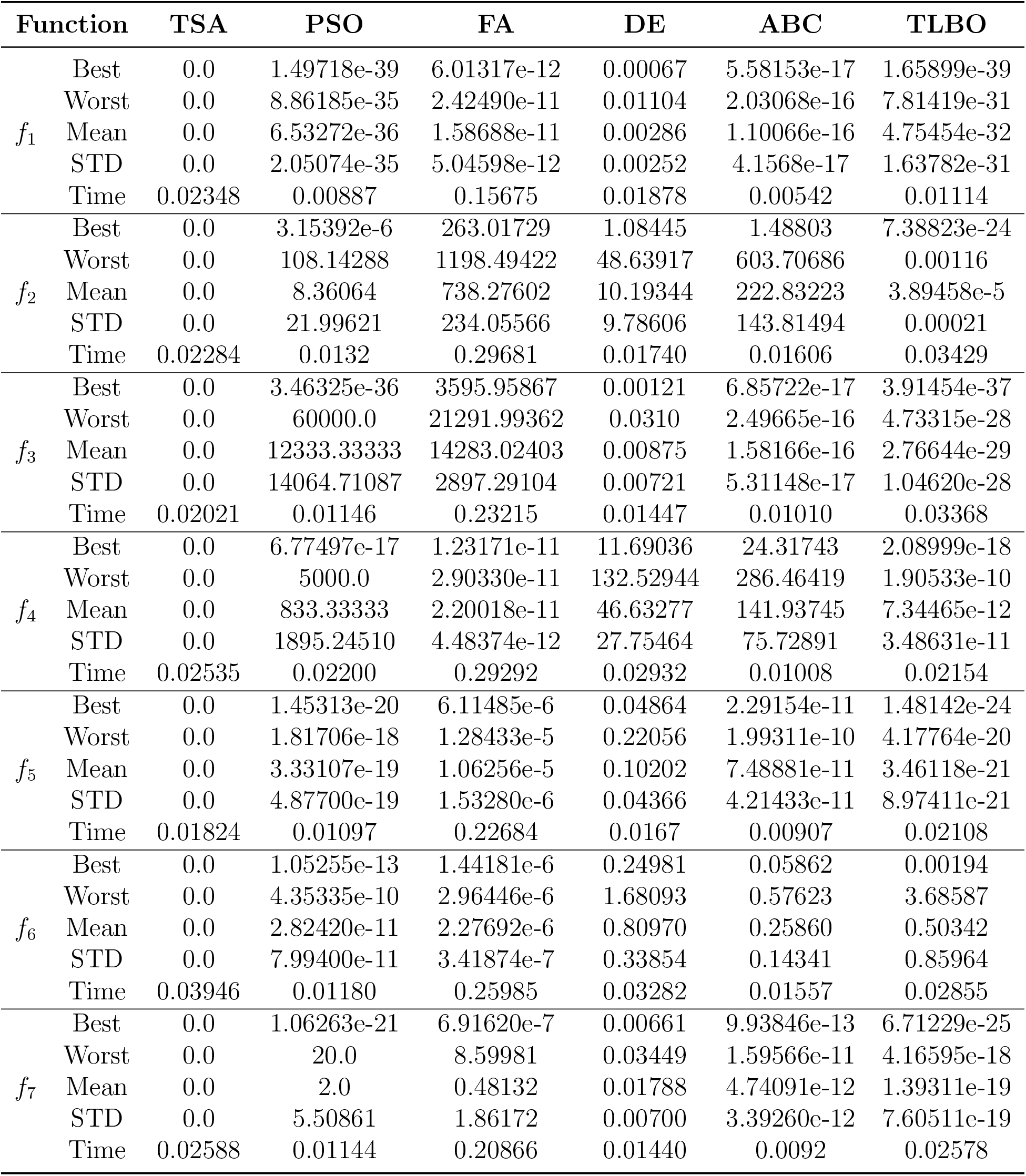
Experimental results of TSA, PSO, FA, DE, ABC, and TLBO algorithms on 14 unimodal functions with dimensionality 10, part I, functions *f*_*i*_ for *i* = 1 … 7.

**Table SI.2:**
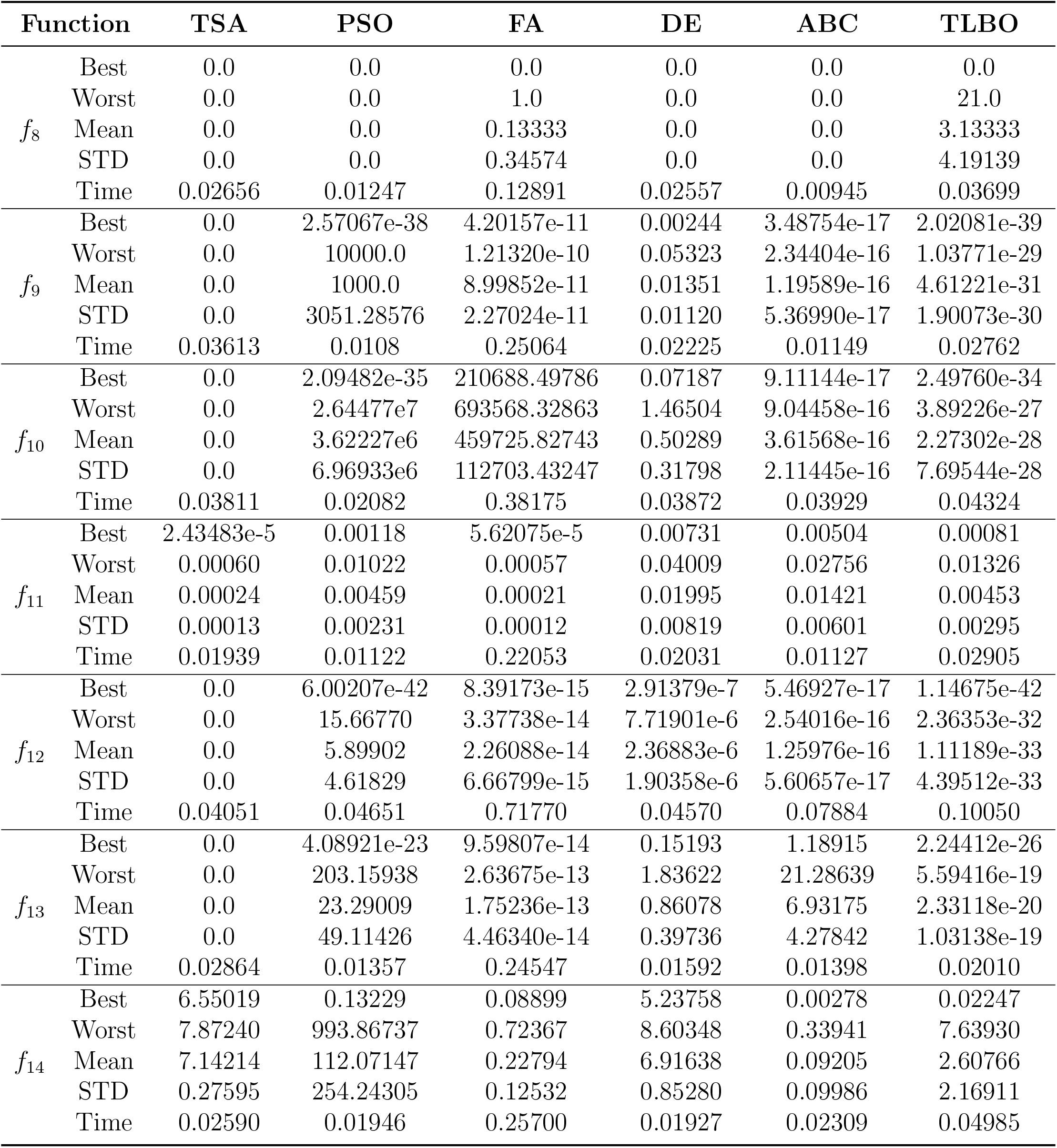
Experimental results of TSA, PSO, FA, DE, ABC, and TLBO algorithms on 14 unimodal functions with dimensionality 10, part II, functions *f*_*i*_ for *i* = 8 … 14.

**Table SI.3:** Experimental results of TSA, PSO, FA, DE, ABC, and TLBO algorithms on 14 unimodal functions with dimensionality 50, part I, functions *f*_*i*_ for *i* = 1 … 7.

| Function |  | TSA | PSO | FA | DE | ABC | TLBO |
| --- | --- | --- | --- | --- | --- | --- | --- |
| $f_1$ | Best | 0.0 | 0.00566 | 7.27283e-10 | 26726.20459 | 0.00121 | 0.80073 |
|  | Worst | 0.0 | 30000.20385 | 1.13847e-9 | 56274.06091 | 0.11855 | 379.04396 |
|  | Mean | 0.0 | 7672.45042 | 9.32253e-10 | 37655.43863 | 0.00959 | 43.16074 |
|  | STD | 0.0 | 8578.54452 | 9.69064e-11 | 7445.17859 | 0.02104 | 83.59772 |
|  | Time | 0.06194 | 0.02691 | 0.40795 | 0.03445 | 0.02319 | 0.05271 |
| $f_2$ | Best | 0.0 | 8083.01221 | 26343.48261 | 0.03532 | 30207.63333 | 850.11809 |
|  | Worst | 0.0 | 29019.53802 | 36812.78455 | 19288.14233 | 42888.91472 | 3372.77999 |
|  | Mean | 0.0 | 19008.13249 | 32403.23317 | 36649.08067 | 36211.38459 | 1919.78994 |
|  | STD | 0.0 | 5237.69666 | 2894.62810 | 26935.76824 | 3164.88770 | 611.29062 |
|  | Time | 0.08624 | 0.05408 | 0.73023 | 5140.99957 | 0.05967 | 0.07186 |
| $f_3$ | Best | 0.0 | 10000.79209 | 110760.12598 | 23317.90803 | 0.0029471 | 33.15641 |
|  | Worst | 0.0 | 200000.00002 | 154896.64960 | 57290.40004 | 68.73835 | 5367.85249 |
|  | Mean | 0.0 | 87001.98657 | 128242.42945 | 40422.10237 | 11.52768 | 641.07483 |
|  | STD | 0.0 | 46912.84064 | 8470.96659 | 8597.24784 | 15.83657 | 1181.74747 |
|  | Time | 0.08712 | 0.03081 | 0.65467 | 0.04135 | 0.03707 | 0.06695 |
| $f_4$ | Best | 0.0 | 22100.38383 | 14095.30150 | 107356.43475 | 42030.46556 | 1532.56205 |
|  | Worst | 0.0 | 113394.11732 | 17120.81244 | 185555.59051 | 67318.63611 | 7627.84490 |
|  | Mean | 0.0 | 51289.49566 | 15631.00522 | 149459.37474 | 54987.91466 | 4421.34759 |
|  | STD | 0.0 | 18590.06180 | 674.26592 | 18657.22168 | 7677.68078 | 1722.58092 |
|  | Time | 0.07776 | 0.03213 | 0.52603 | 0.04129 | 0.03056 | 0.05662 |
| $f_5$ | Best | 0.0 | 5.22466 | 39.69779 | 765.04807 | 0.30031 | 4.81668 |
|  | Worst | 0.0 | 600.96472 | 66.16936 | 1255.20281 | 2.55359 | 73.21633 |
|  | Mean | 0.0 | 293.16944 | 50.95405 | 1007.26907 | 0.84523 | 29.29733 |
|  | STD | 0.0 | 124.45490 | 6.24908 | 124.82628 | 0.53657 | 18.82647 |
|  | Time | 0.08333 | 0.03650 | 0.63219 | 0.03455 | 0.03428 | 0.07993 |
| $f_6$ | Best | 0.0 | 25.98913 | 52.18881 | 88.35429 | 57.33431 | 17.79045 |
|  | Worst | 1.42108e-14 | 47.19694 | 58.03175 | 96.12514 | 73.93299 | 33.50344 |
|  | Mean | 8.52651e-15 | 37.60352 | 54.47123 | 92.63260 | 65.65450 | 25.07611 |
|  | STD | 7.08088e-15 | 5.95083 | 1.42114 | 1.92489 | 4.06774 | 3.49861 |
|  | Time | 0.06501 | 0.04046 | 0.65660 | 0.02542 | 0.02188 | 0.06302 |
| $f_7$ | Best | 0.0 | 20.30314 | 156.45757 | 89.07719 | 0.02741 | 0.22032 |
|  | Worst | 0.0 | 120.04493 | 201.42351 | 1.53366e12 | 0.14100 | 8.52087 |
|  | Mean | 0.0 | 65.96133 | 179.31771 | 5.12533e10 | 0.05723 | 3.48042 |
|  | STD | 0.0 | 22.39311 | 9.50055 | 2.799823e11 | 0.02585 | 2.12310 |
|  | Time | 0.08020 | 0.02636 | 0.48331 | 0.03449 | 0.03979 | 0.05623 |

**Table SI.4:**
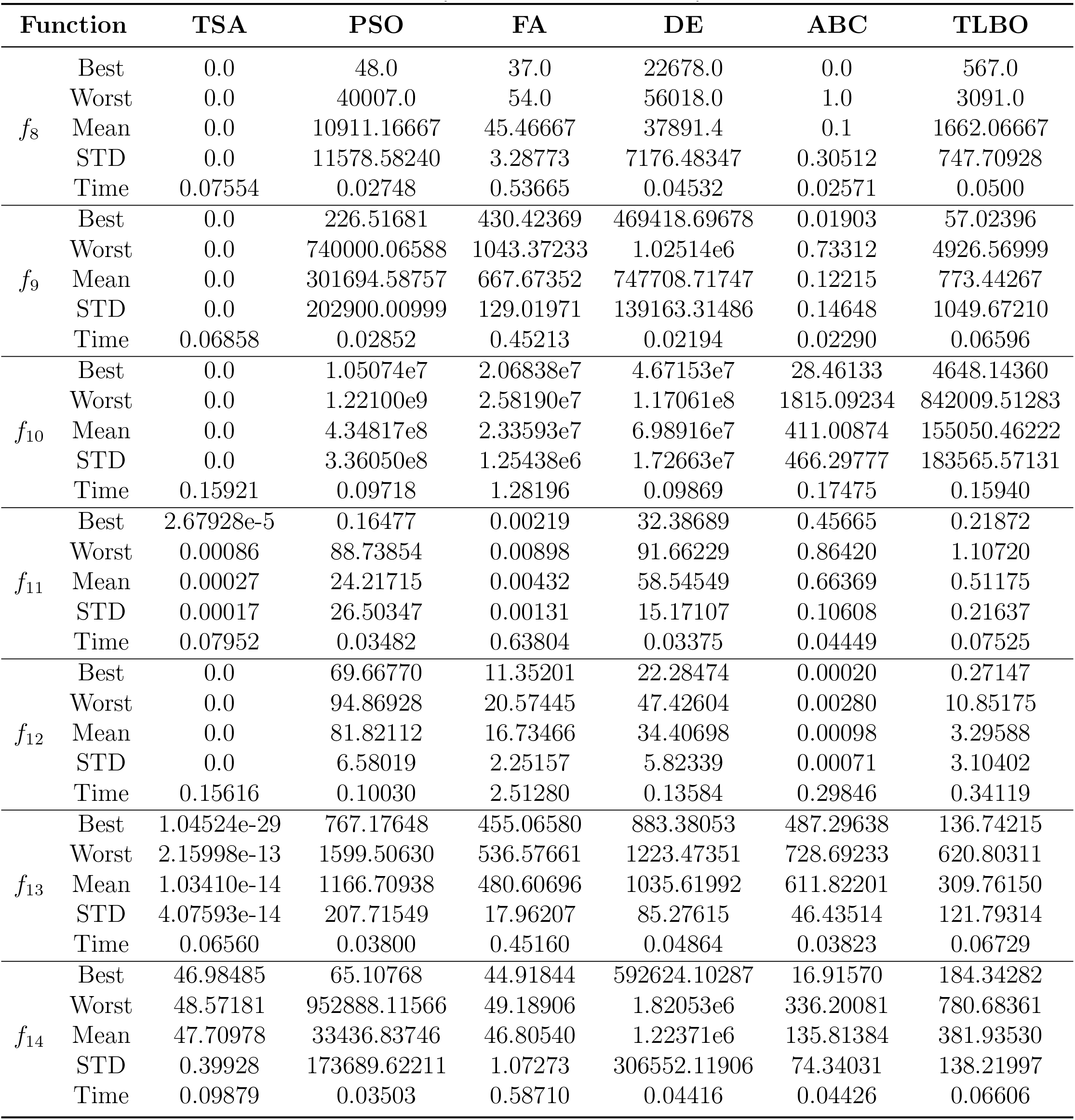
Experimental results of TSA, PSO, FA, DE, ABC, and TLBO algorithms on 14 unimodal functions with dimensionality 50, part II, functions *f*_*i*_ for *i* = 8 … 14.

**Table SI.5:**
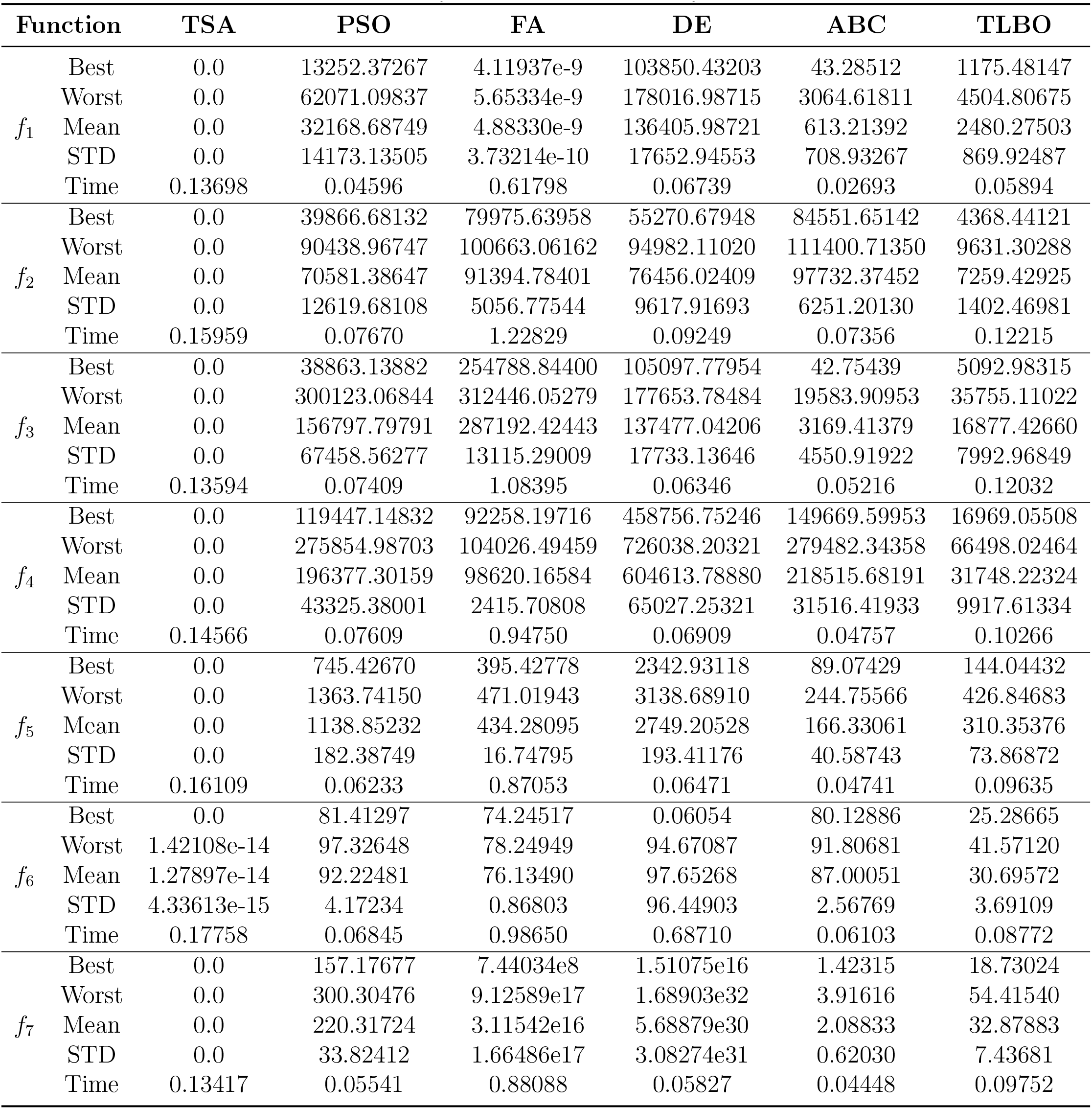
Experimental results of TSA, PSO, FA, DE, ABC, and TLBO algorithms on 14 unimodal functions with dimensionality 100, part I, functions *f*_*i*_ for *i* = 1 … 7.

**Table SI.6:**
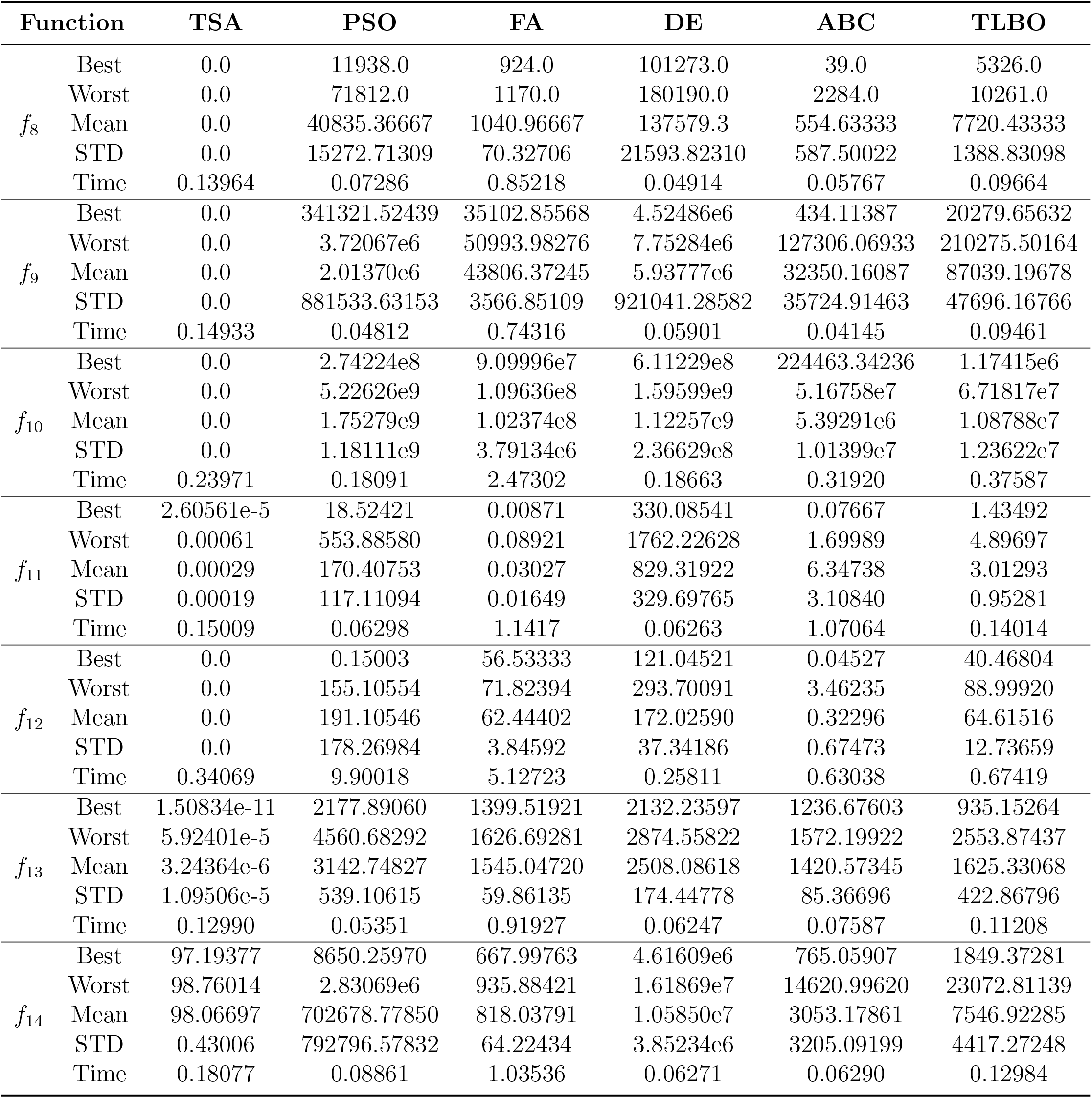
Experimental results of TSA, PSO, FA, DE, ABC, and TLBO algorithms on 14 unimodal functions with dimensionality 100, part II, functions *f*_*i*_ for *i* = 8 … 14.

**Table SI.7:**
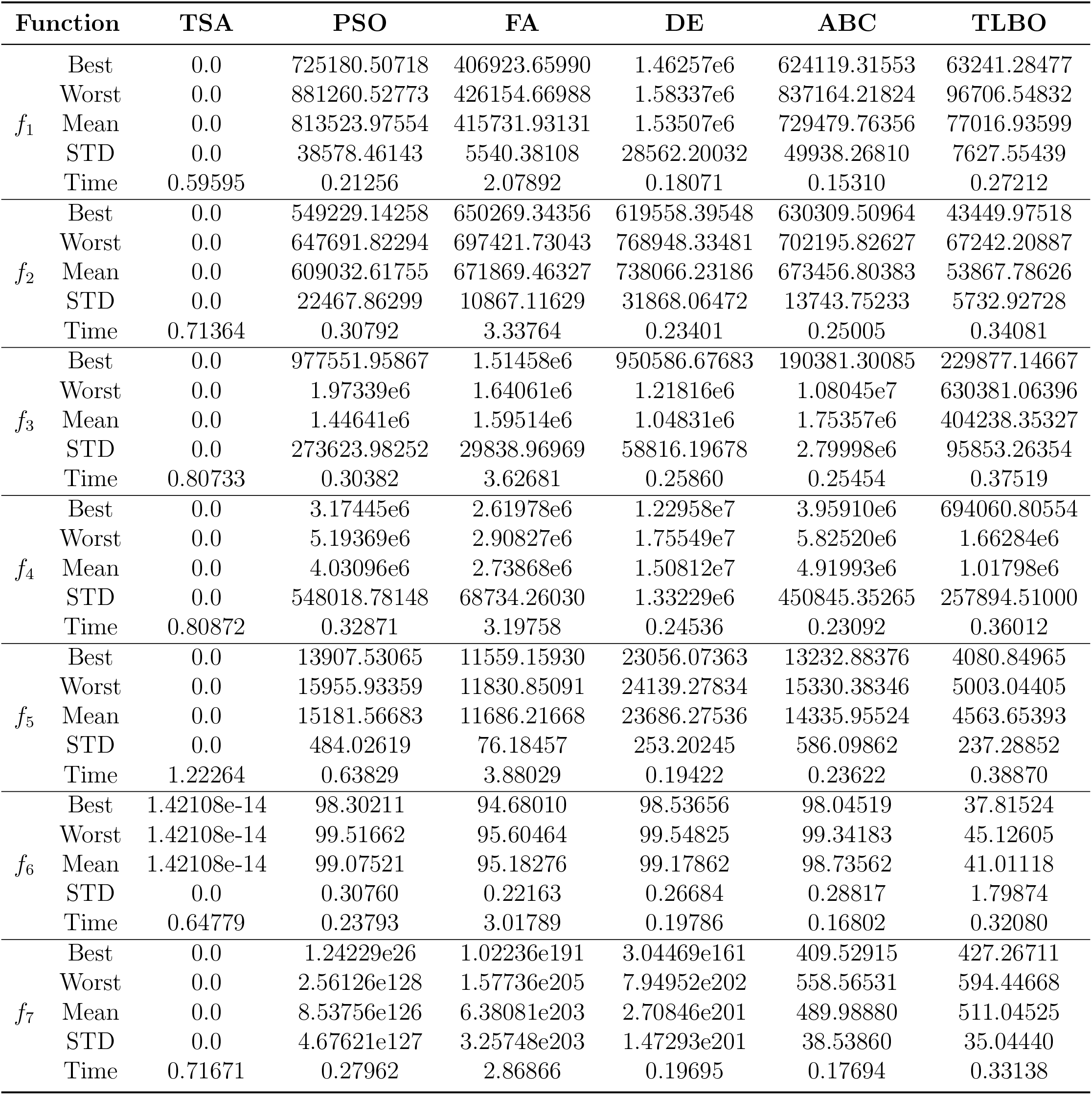
Experimental results of TSA, PSO, FA, DE, ABC, and TLBO algorithms on 14 unimodal functions with dimensionality 500, part I, functions *f*_*i*_ for *i* = 1 … 7.

**Table SI.8:** Experimental results of TSA, PSO, FA, DE, ABC, and TLBO algorithms on 14 unimodal functions with dimensionality 500, part II, functions *f*_*i*_ for *i* = 8 … 14.

| Function |  | TSA | PSO | FA | DE | ABC | TLBO |
| --- | --- | --- | --- | --- | --- | --- | --- |
| $f_8$ | Best | 0.0 | 740435.0 | 481314.0 | 1.46261e6 | 655191.0 | 85977.0 |
|  | Worst | 0.0 | 902287.0 | 516380.0 | 1.58402e6 | 814652.0 | 129428.0 |
|  | Mean | 0.0 | 826535.43333 | 499195.06667 | 1.53537e6 | 734925.23333 | 100020.36667 |
|  | STD | 0.0 | 39930.42562 | 8122.14444 | 28568.86615 | 43161.83623 | 8645.09009 |
|  | Time | 0.61643 | 0.23663 | 2.64437 | 0.18758 | 0.16643 | 0.36080 |
| $f_9$ | Best | 0.0 | 1.65502e8 | 7.69170e7 | 3.56909e8 | 1.45974e8 | 1.27146e7 |
|  | Worst | 0.0 | 2.03505e8 | 8.55635e7 | 3.92615e8 | 2.04450e8 | 2.00315e7 |
|  | Mean | 0.0 | 1.87459e8 | 8.30437e7 | 3.78555e8 | 1.83009e8 | 1.71554e7 |
|  | STD | 0.0 | 1.09836e7 | 1.83808e6 | 7.91620e6 | 1.38803e7 | 1.82643e6 |
|  | Time | 0.59004 | 0.24976 | 2.29803 | 0.15865 | 0.17556 | 0.27879 |
| $f_{10}$ | Best | 0.0 | 1.58113e10 | 4.00707e9 | 3.51121e10 | 2.96068e10 | 9.36988e8 |
|  | Worst | 0.0 | 4.28044e10 | 4.38831e9 | 5.90791e10 | 5.26760e10 | 2.32025e9 |
|  | Mean | 0.0 | 2.51279e10 | 4.18209e9 | 4.30779e10 | 4.20229e10 | 1.64839e9 |
|  | STD | 0.0 | 5.70366e9 | 9.72621e7 | 6.14113e9 | 5.66690e9 | 3.38430e8 |
|  | Time | 1.22110 | 0.91098 | 10.33665 | 0.84552 | 1.53772 | 1.73280 |
| $f_{11}$ | Best | 4.85599e-5 | 18333.15676 | 6179.21665 | 53361.63195 | 16804.36571 | 143.33145 |
|  | Worst | 0.00106 | 27009.59654 | 6999.78733 | 61218.38881 | 26984.13609 | 324.56579 |
|  | Mean | 0.00031 | 23082.32631 | 6580.32298 | 58273.46197 | 22858.39268 | 235.38178 |
|  | STD | 0.00023 | 2170.40935 | 158.55609 | 1857.12175 | 2208.78435 | 48.04955 |
|  | Time | 0.74848 | 0.34378 | 3.86230 | 0.28002 | 0.34380 | 0.47425 |
| $f_{12}$ | Best | 0.0 | 962.49971 | 5582.53299 | 356887.94922 | 4089.15752 | 3537.94759 |
|  | Worst | 0.0 | 986.13482 | 7298.47475 | 3.31423e6 | 146793.56898 | 5641.80956 |
|  | Mean | 0.0 | 974.10952 | 6328.32474 | 1.51729e6 | 56734.37776 | 4407.38716 |
|  | STD | 0.0 | 6.37547 | 382.40563 | 876860.87038 | 47154.75458 | 430.81666 |
|  | Time | 1.44452 | 0.69272 | 18.90528 | 1.30118 | 2.81625 | 3.04662 |
| $f_{13}$ | Best | 0.00797 | 19557.31864 | 185.52719 | 62420.38153 | 8391.84933 | 10469.41456 |
|  | Worst | 9.06537 | 23880.27457 | 8744.30549 | 34897.47245 | 10249.62214 | 21490.92463 |
|  | Mean | 0.98107 | 21696.56739 | 9144.10422 | 319950.41580 | 9137.54108 | 14205.78842 |
|  | STD | 1.86606 | 1304.94735 | 8429.20196 | 13296.04502 | 414.98910 | 2257.10354 |
|  | Time | 0.70937 | 0.28206 | 3.10761 | 0.20351 | 0.25163 | 0.36405 |
| $f_{14}$ | Best | 498.16276 | 2.94243e7 | 1.20733e7 | 8.40511e7 | 2.88270e7 | 338867.21201 |
|  | Worst | 498.69215 | 4.31867e7 | 1.36132e7 | 9.35240e7 | 4.01719e7 | 596766.89085 |
|  | Mean | 498.41361 | 3.78163e7 | 1.28485e7 | 8.98129e7 | 3.35809e7 | 444367.46024 |
|  | STD | 0.15010 | 3.13191e6 | 389423.33613 | 2.57019e6 | 3.16303e6 | 64500.50536 |
|  | Time | 0.69365 | 0.29930 | 2.73897 | 0.24004 | 0.23114 | 0.36428 |

### SI.2 Results for Multimodel Benchmark Functions

**Table SI.9:**
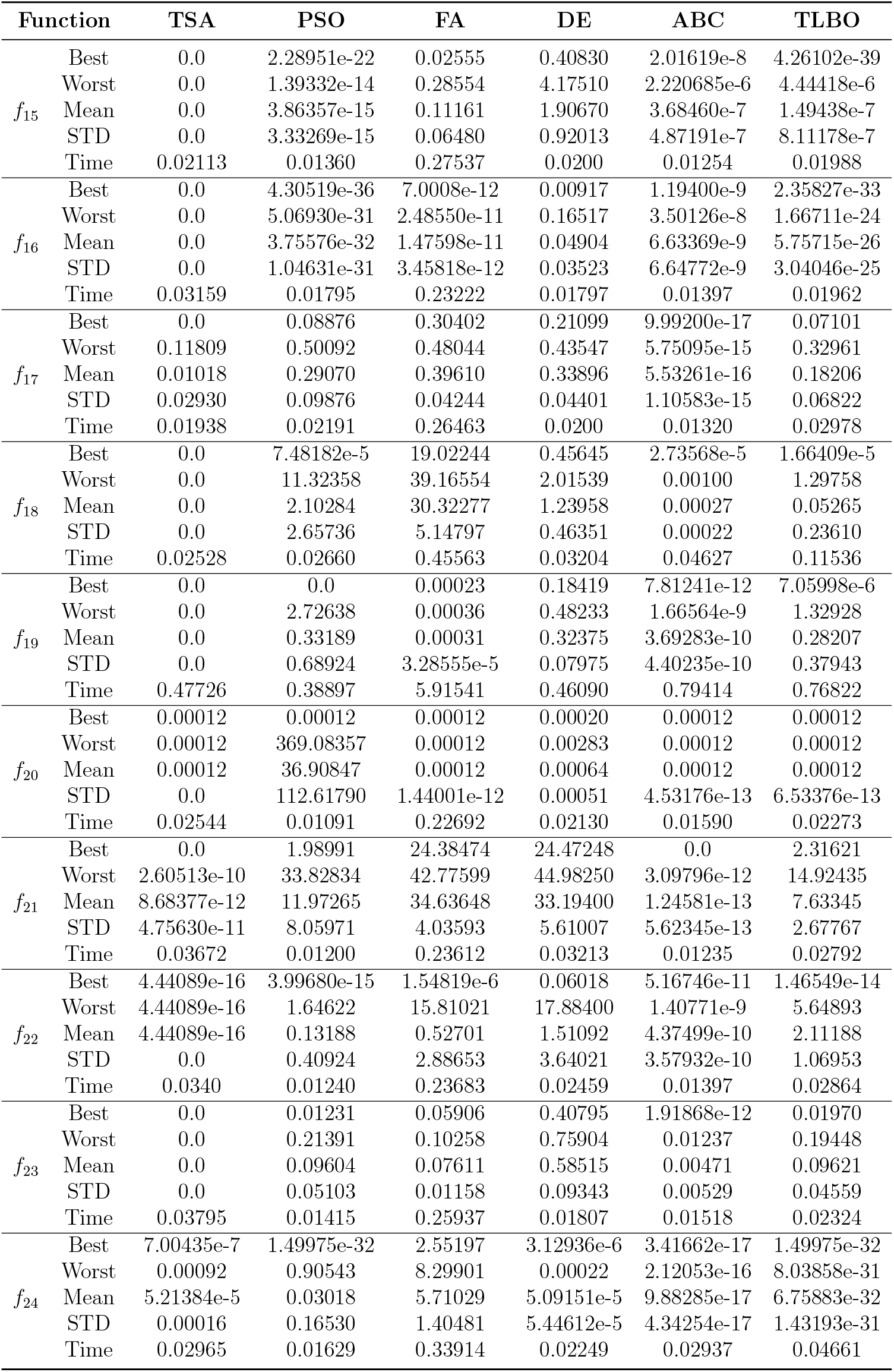
Experimental results of TSA, PSO, FA, DE, ABC, and TLBO algorithms on 10 multimodal functions with dimensionality 10.

**Table SI.10:**
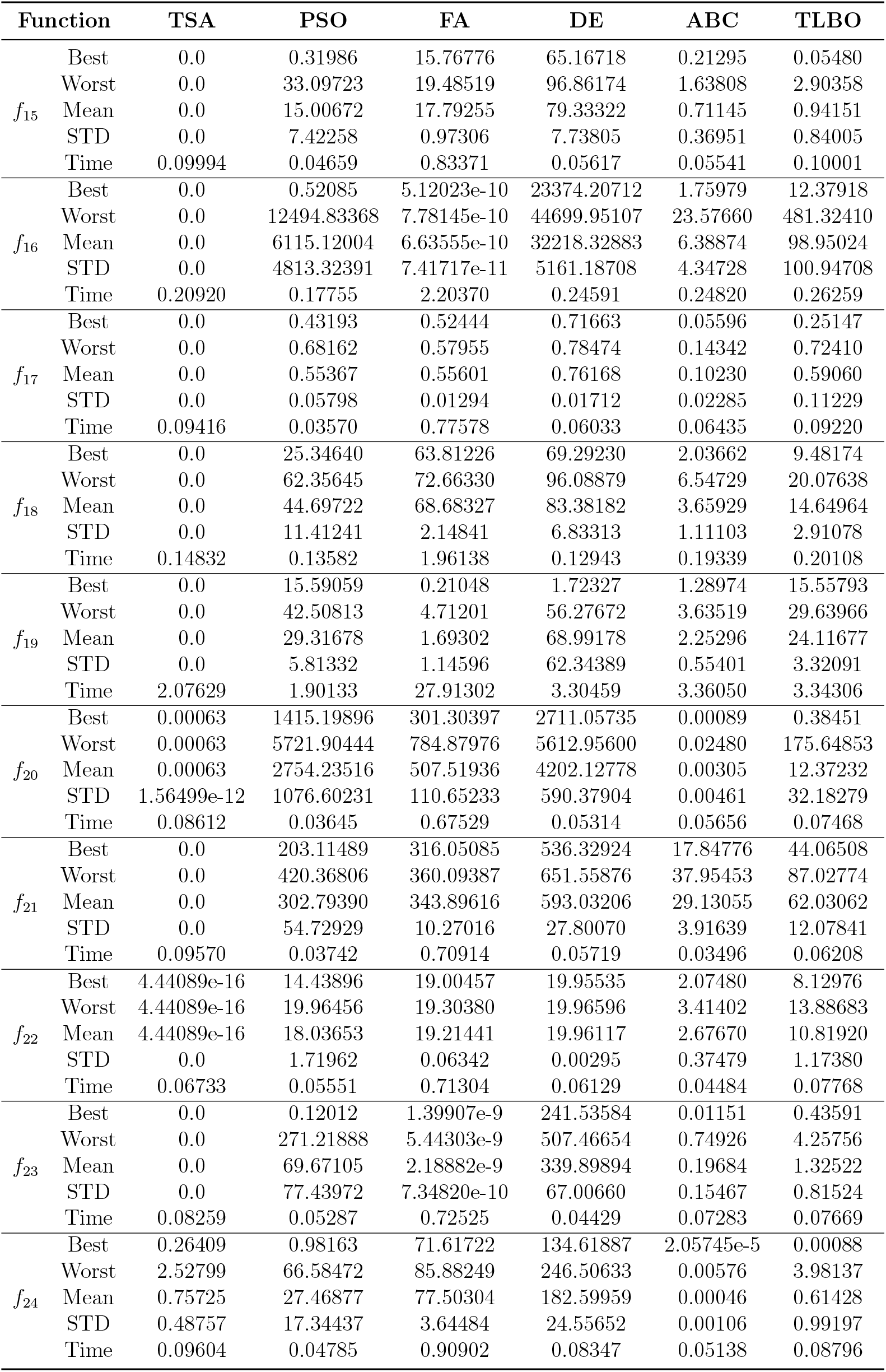
Experimental results of TSA, PSO, FA, DE, ABC, and TLBO algorithms on 10 multimodal functions with dimensionality 50.

**Table SI.11:**
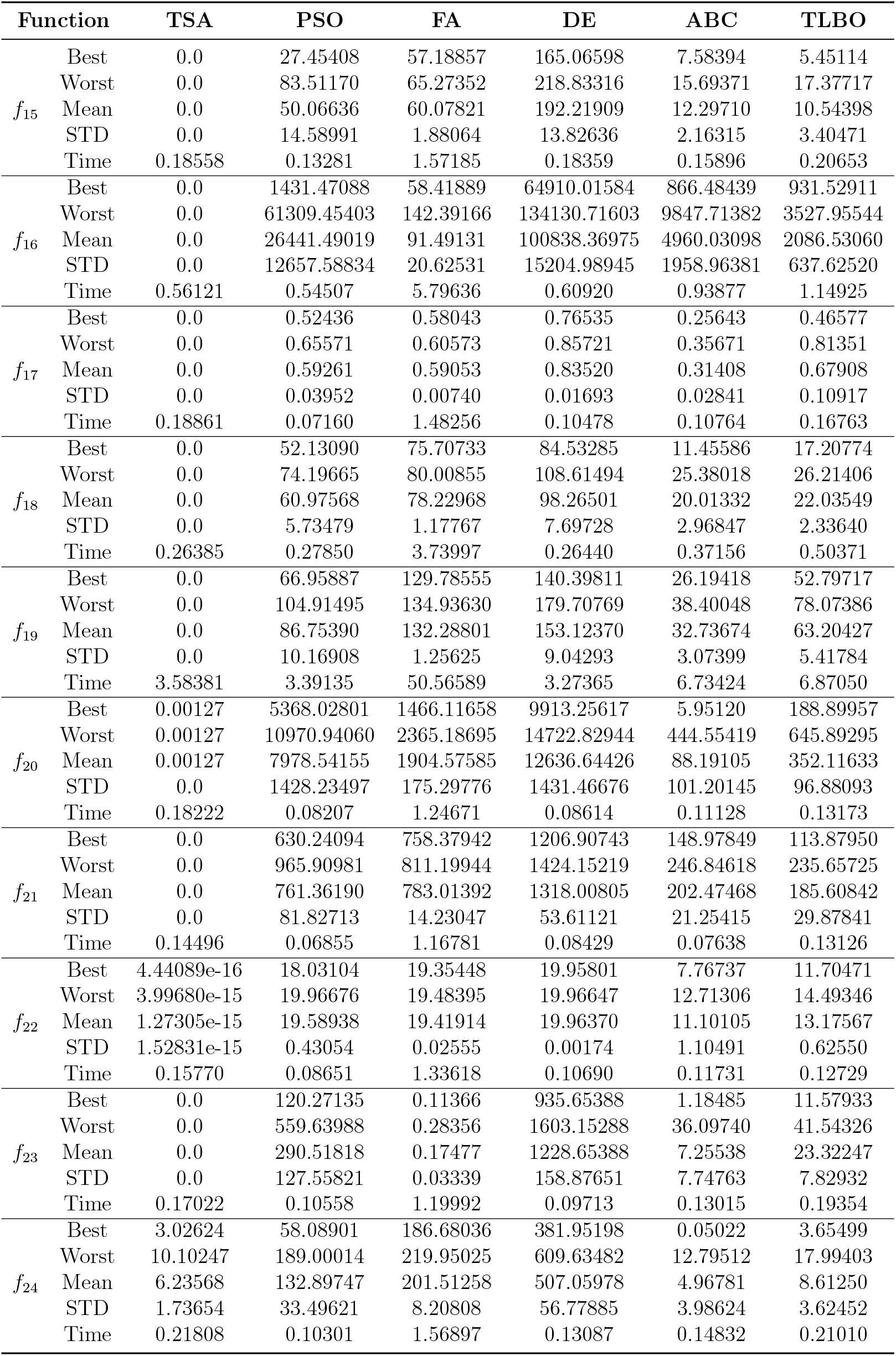
Experimental results of TSA, PSO, FA, DE, ABC, and TLBO algorithms on 10 multimodal functions with dimensionality 100.

**Table SI.12:** Experimental results of TSA, PSO, FA, DE, ABC, and TLBO algorithms on 10 multimodal functions with dimensionality 100.

| Function | TSA | PSO | FA | DE | ABC | TLBO |  |
| --- | --- | --- | --- | --- | --- | --- | --- |
| $f_{15}$ | Best | 0.0 | 685.02532 | 697.43691 | 1074.95201 | 628.36392 | 241.61472 |
|  | Worst | 0.0 | 849.91662 | 717.22485 | 1410.68244 | 716.06335 | 295.35531 |
|  | Mean | 0.0 | 755.84453 | 706.69010 | 1215.60356 | 663.67673 | 267.47664 |
|  | STD | 0.0 | 39.38074 | 6.16776 | 99.51038 | 24.14594 | 17.04008 |
|  | Time | 0.77271 | 0.39921 | 4.23752 | 0.49104 | 0.63060 | 0.53377 |
| $f_{16}$ | Best | 0.0 | 504830.17749 | 316861.14211 | 557425.23109 | 502396.13477 | 47396.81599 |
|  | Worst | 0.0 | 620725.35638 | 330049.58351 | 784794.78941 | 623870.56457 | 100234.11671 |
|  | Mean | 0.0 | 558142.94850 | 323114.85610 | 667929.33820 | 556408.91819 | 69834.31367 |
|  | STD | 0.0 | 29026.90408 | 3452.83597 | 57002.36795 | 28187.77110 | 10086.57514 |
|  | Time | 9.48841 | 11.63966 | 104.90662 | 11.47786 | 17.30530 | 17.37130 |
| $f_{17}$ | Best | 0.0 | 0.70594 | 0.67066 | 0.91442 | 0.70183 | 0.77788 |
|  | Worst | 0.0 | 0.79334 | 0.68085 | 0.93721 | 0.74497 | 0.91694 |
|  | Mean | 0.0 | 0.73521 | 0.67464 | 0.92879 | 0.72964 | 0.86066 |
|  | STD | 0.0 | 0.02050 | 0.00238 | 0.00495 | 0.01119 | 0.04240 |
|  | Time | 0.81405 | 0.41883 | 4.74100 | 0.42357 | 0.76999 | 1.09397 |
| $f_{18}$ | Best | 0.0 | 91.50574 | 100.55886 | 104.45396 | 76.87046 | 28.07302 |
|  | Worst | 0.0 | 107.47388 | 102.73515 | 122.60910 | 89.50815 | 36.53280 |
|  | Mean | 0.0 | 99.65688 | 101.52351 | 113.49883 | 82.14496 | 32.19929 |
|  | STD | 0.0 | 3.71022 | 0.54016 | 4.34074 | 3.61142 | 2.16806 |
|  | Time | 1.30685 | 1.24709 | 16.11589 | 1.18098 | 2.18229 | 2.22547 |
| $f_{19}$ | Best | 0.0 | 707.17436 | 838.13117 | 946.10139 | 651.28153 | 407.72498 |
|  | Worst | 0.0 | 814.62554 | 856.23496 | 976.09523 | 730.99591 | 456.54569 |
|  | Mean | 0.0 | 747.24513 | 844.95474 | 963.30559 | 694.45546 | 429.48022 |
|  | STD | 0.0 | 24.71094 | 4.25315 | 7.18941 | 20.23220 | 12.05624 |
|  | Time | 14.64775 | 14.01315 | 205.68801 | 13.38510 | 27.90171 | 27.99851 |
| $f_{20}$ | Best | 0.00636 | 72903.90437 | 51072.90337 | 74303.27553 | 61161.29566 | 9093.62017 |
|  | Worst | 0.00636 | 87162.32702 | 54368.88901 | 116869.34434 | 75887.95755 | 16219.42905 |
|  | Mean | 0.00636 | 80304.72445 | 52658.19358 | 91245.78769 | 68652.98737 | 12235.94100 |
|  | STD | 1.45016e-11 | 3674.26605 | 731.60206 | 9156.62267 | 3873.36467 | 1909.84856 |
|  | Time | 0.76201 | 0.43561 | 4.24607 | 0.42216 | 0.69004 | 0.60276 |
| $f_{21}$ | Best | 0.0 | 5778.91603 | 5632.81275 | 7082.93685 | 4652.20213 | 2521.04619 |
|  | Worst | 0.0 | 6390.32631 | 5747.90485 | 8049.08764 | 5654.23349 | 3025.43478 |
|  | Mean | 0.0 | 6070.43746 | 5683.73594 | 7480.47112 | 5144.80007 | 2799.21265 |
|  | STD | 0.0 | 131.48467 | 35.84680 | 239.33614 | 217.69202 | 132.34141 |
|  | Time | 0.72142 | 0.35920 | 3.47144 | 0.27871 | 0.44006 | 0.47190 |
| $f_{22}$ | Best | 4.44089e-16 | 19.96413 | 20.18037 | 19.96592 | 20.02075 | 15.00832 |
|  | Worst | 3.99680e-15 | 20.40762 | 20.21873 | 19.97219 | 20.21104 | 17.19914 |
|  | Mean | 3.75995e-15 | 19.98051 | 20.19349 | 19.96690 | 20.12779 | 16.19672 |
|  | STD | 9.01352e-16 | 0.08067 | 0.00910 | 0.00135 | 0.04857 | 0.54794 |
|  | Time | 0.73736 | 0.42446 | 4.12196 | 0.35824 | 0.54280 | 0.53293 |
| $f_{23}$ | Best | 0.0 | 6527.62456 | 3663.31293 | 13164.17209 | 5621.27951 | 570.17156 |
|  | Worst | 0.0 | 7932.34474 | 3836.39202 | 14251.33198 | 7535.47796 | 871.35893 |
|  | Mean | 0.0 | 7322.71577 | 3742.58738 | 13816.63067 | 6554.93957 | 694.15242 |
|  | STD | 0.0 | 347.20615 | 49.86342 | 257.05980 | 454.39648 | 68.64798 |
|  | Time | 0.84823 | 0.47093 | 4.89903 | 0.43681 | 0.63859 | 0.73341 |
| $f_{24}$ | Best | 120.31851 | 1996.08386 | 1964.51763 | 4748.33112 | 1617.75422 | 211.83756 |
|  | Worst | 159.45170 | 2586.43175 | 2067.38091 | 6005.64353 | 2146.82124 | 330.13186 |
|  | Mean | 136.56759 | 2305.18196 | 2017.57661 | 5680.56274 | 1958.82025 | 255.63458 |
|  | STD | 10.12540 | 145.29707 | 27.25009 | 233.78997 | 113.90446 | 32.11593 |
|  | Time | 0.88556 | 0.38478 | 4.36236 | 0.39394 | 0.62064 | 0.58709 |

### SI.3 Convergence Curves for Unimodal Benchmark Functions

**Figure SI.1:**
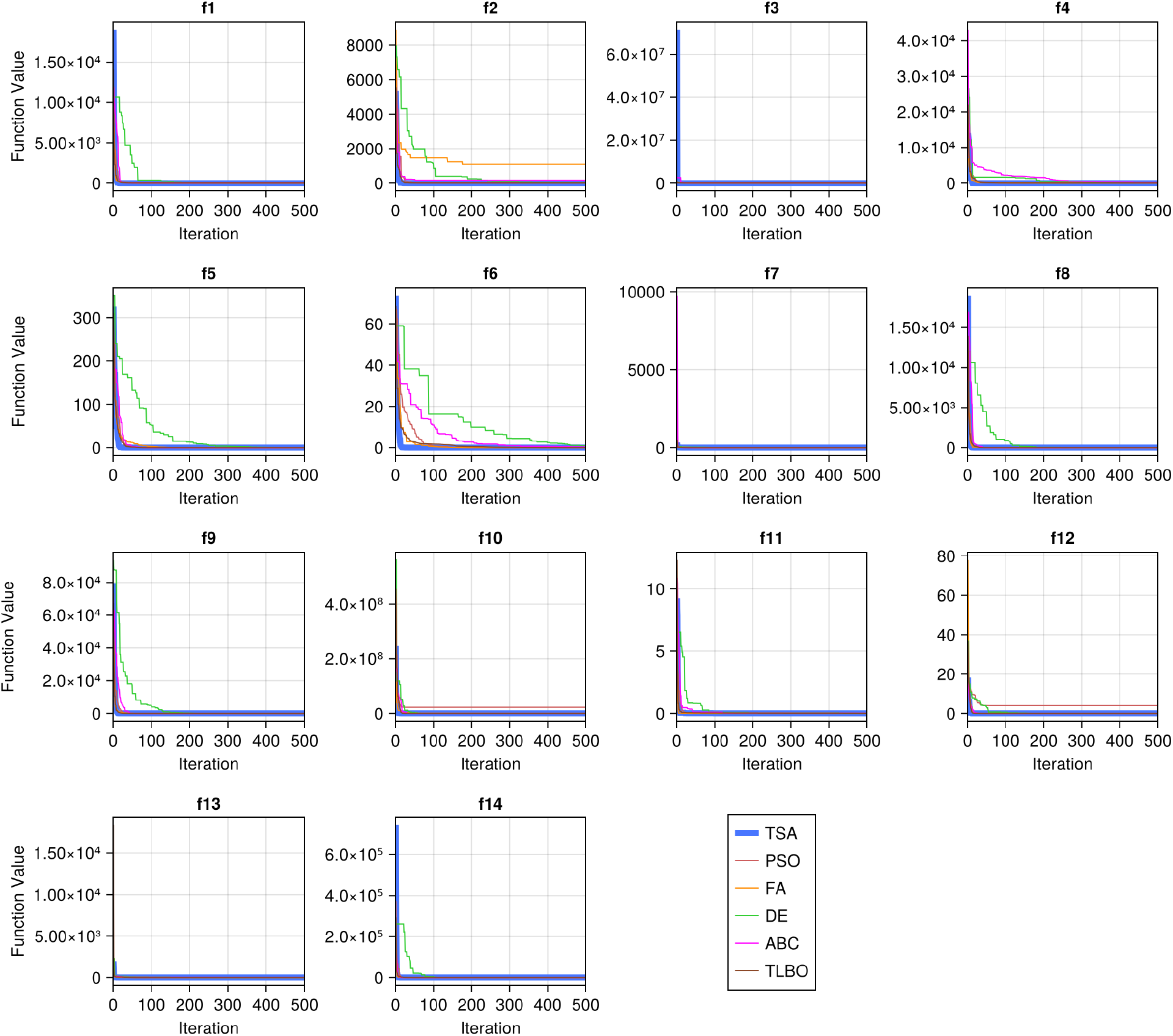
Convergence curves of TSA, PSO, FA, DE, ABC, and TLBO algorithms on 14 unimodal functions with dimensionality 10.

**Figure SI.2:**
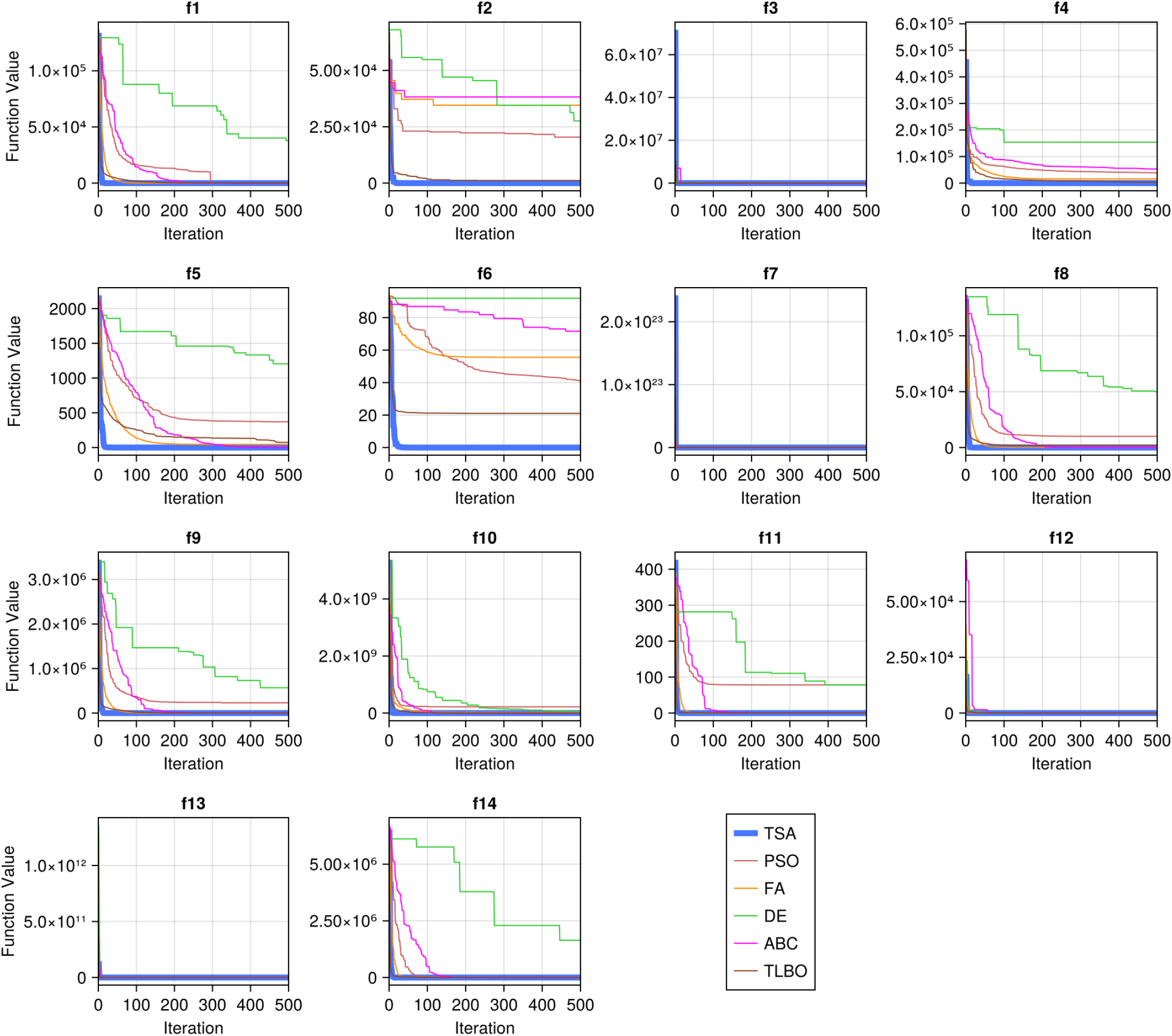
Convergence curves of TSA, PSO, FA, DE, ABC, and TLBO algorithms on 14 unimodal functions with dimensionality 50.

**Figure SI.3:**
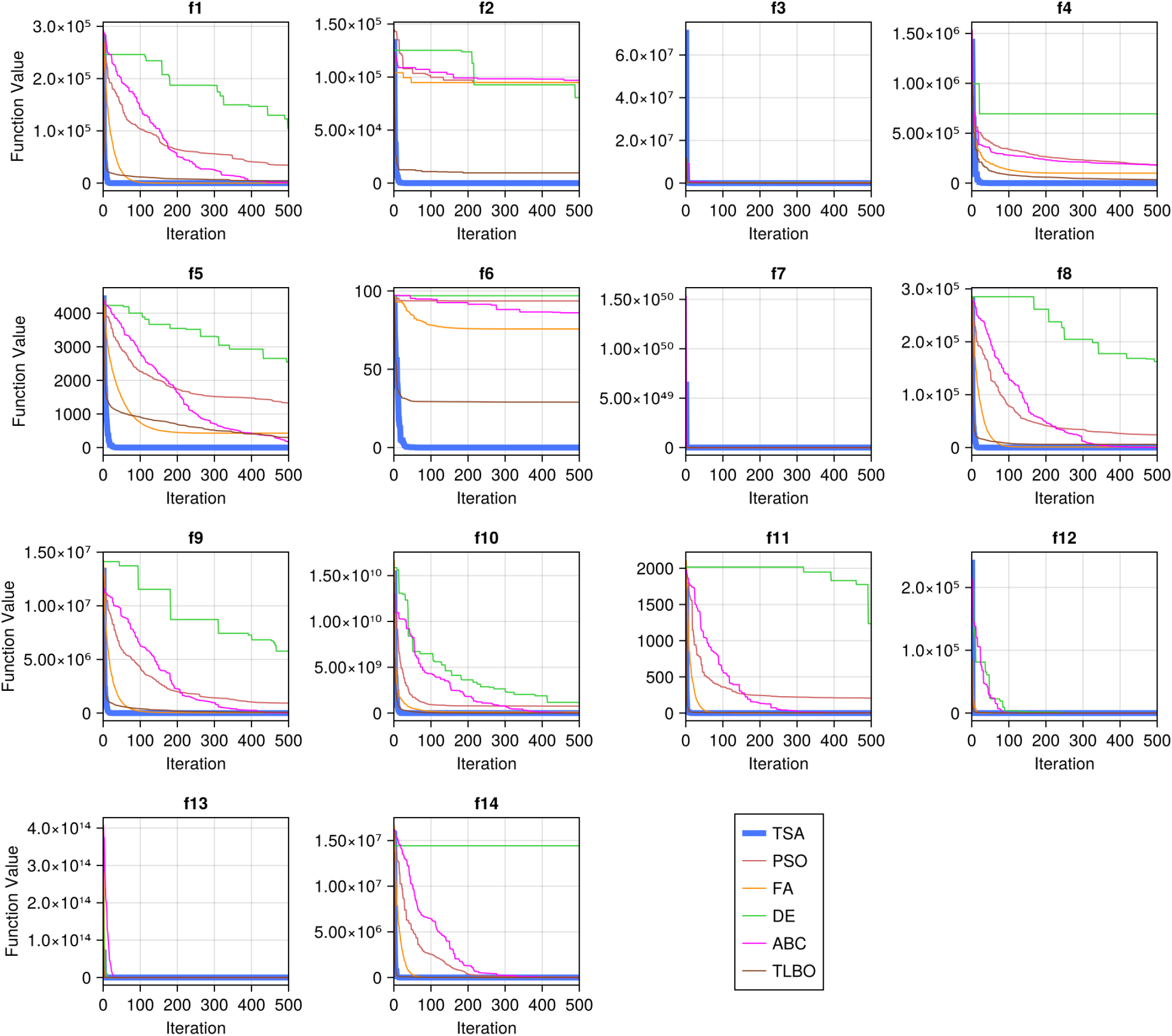
Convergence curves of TSA, PSO, FA, DE, ABC, and TLBO algorithms on 14 unimodal functions with dimensionality 100.

**Figure SI.4:**
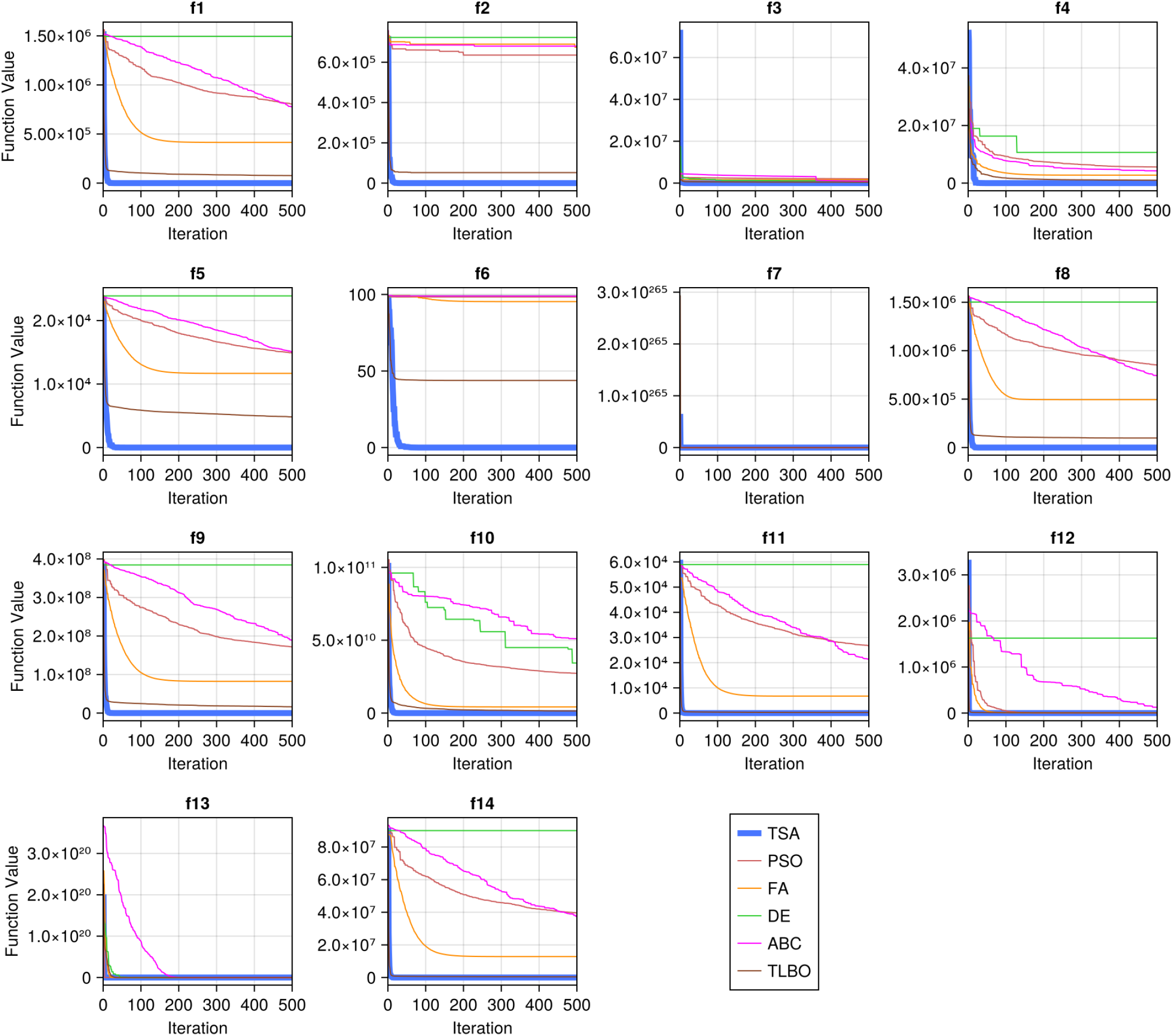
Convergence curves of TSA, PSO, FA, DE, ABC, and TLBO algorithms on 14 unimodal functions with dimensionality 500.

### SI.4 Convergence Curves for Multimodal Benchmark Functions

**Figure SI.5:**
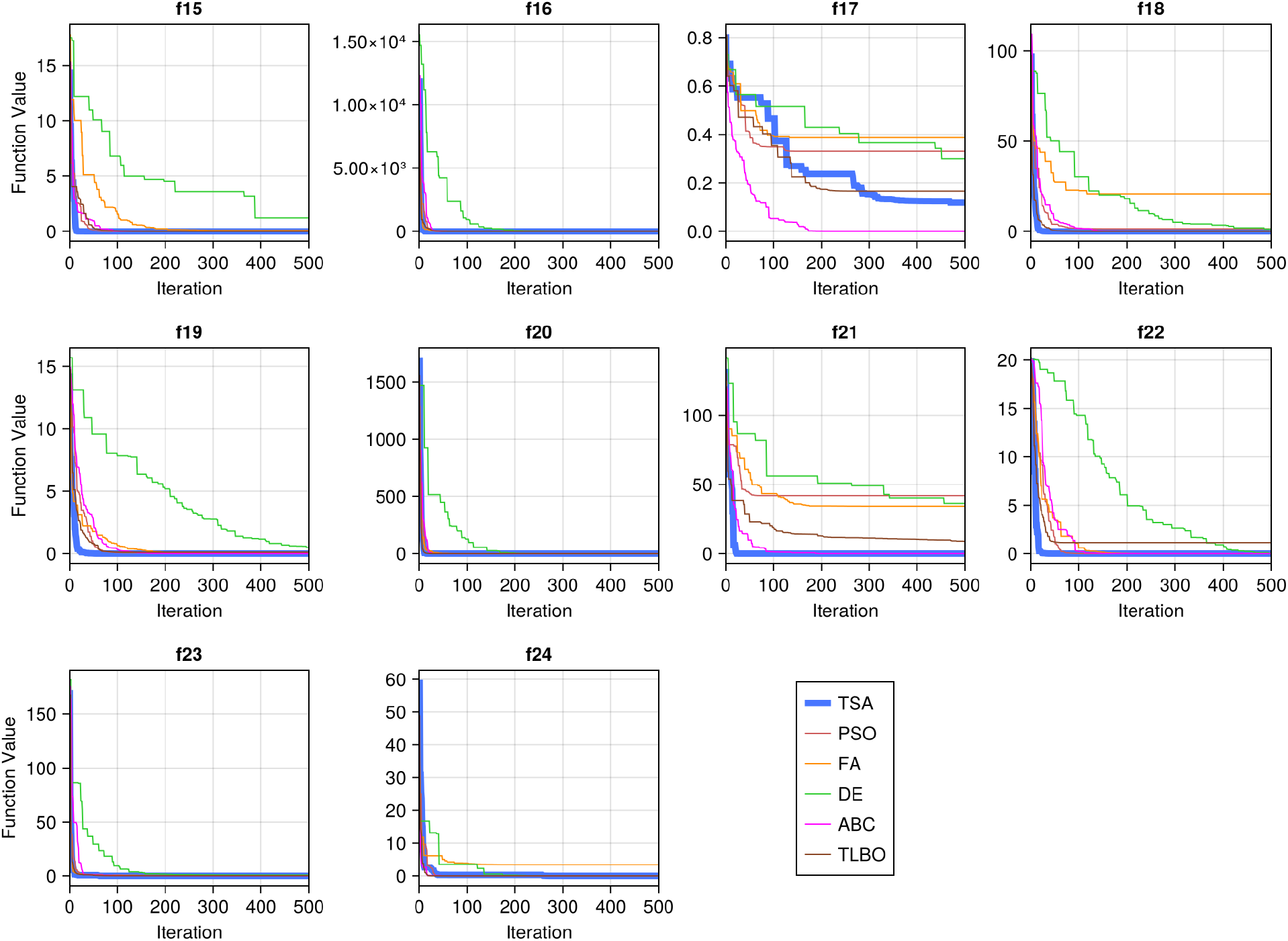
Convergence curves of TSA, PSO, FA, DE, ABC, and TLBO algorithms on 10 multimodal functions with dimensionality 10.

**Figure SI.6:**
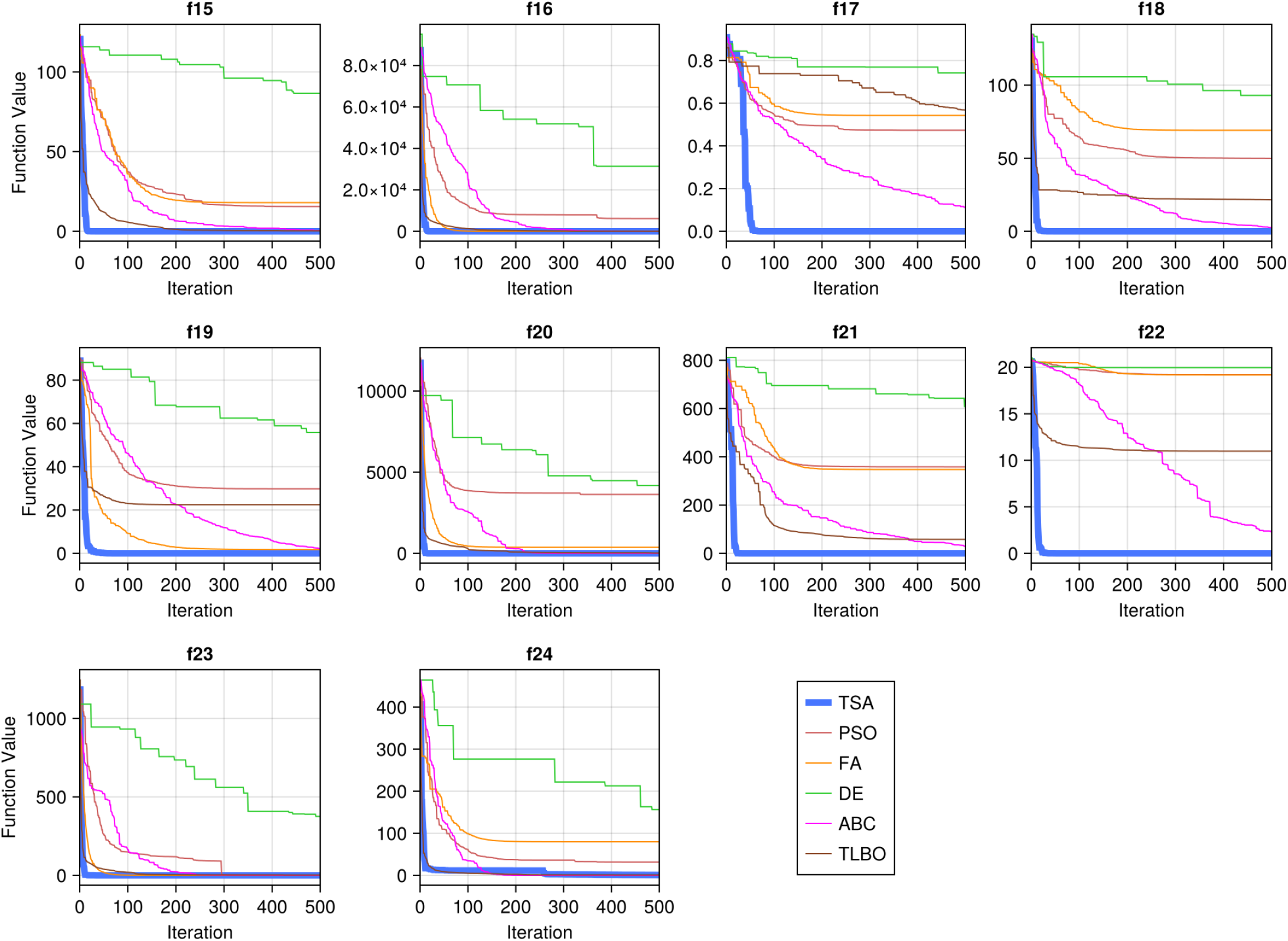
Convergence curves of TSA, PSO, FA, DE, ABC, and TLBO algorithms on 10 multimodal functions with dimensionality 50.

**Figure SI.7:**
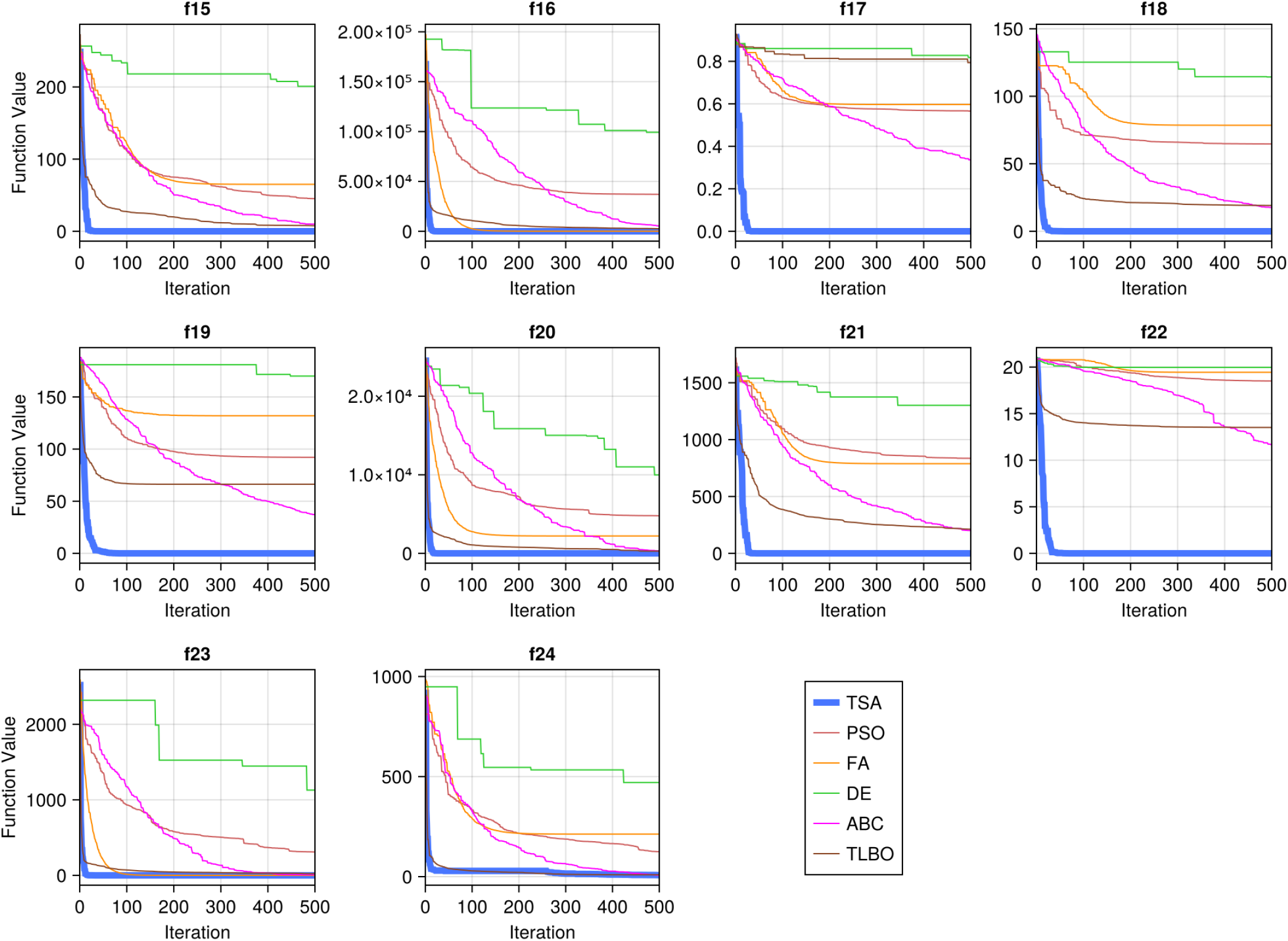
Convergence curves of TSA, PSO, FA, DE, ABC, and TLBO algorithms on 10 multimodal functions with dimensionality 100.

**Figure SI.8:**
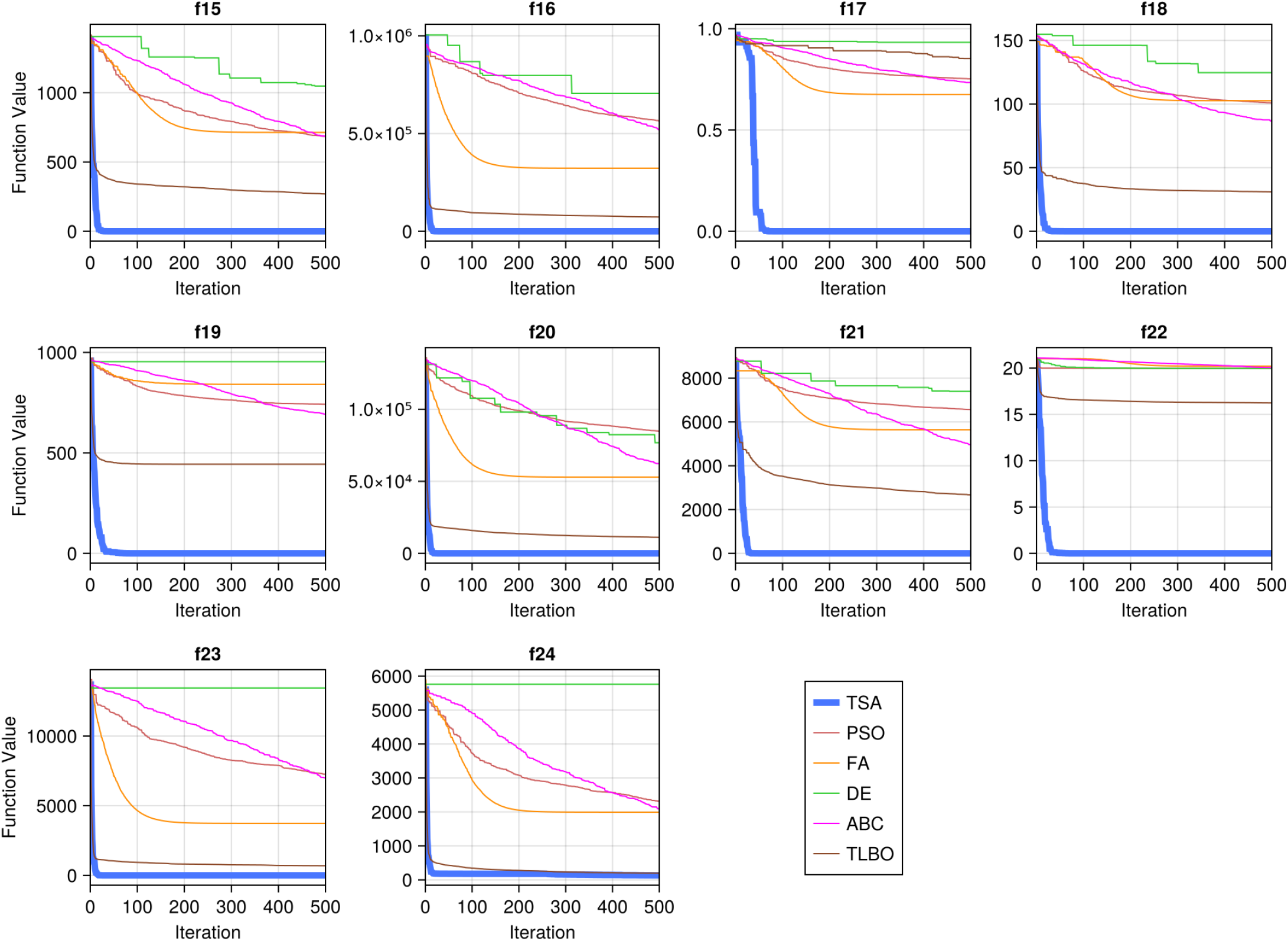
Convergence curves of TSA, PSO, FA, DE, ABC, and TLBO algorithms on 10 multimodal functions with dimensionality 500.

### SI.5 Results of Parameter Estimation

**Table SI.13:** LLSEs and computation time for the LK chemotherapy model parameter estimation using the synthetic data for population sizes *N*_*p*_ = 5, 10, 15, and 20.

| $N_p$ | | **TSA** | **PSO** | **FA** | **DE** | **ABC** | **TLBO** |
| --- | --- | --- | --- | --- | --- | --- | --- |
| 5 | Best | 0.00831 | 0.02923 | 0.36897 | 0.01166 | 0.10131 | 0.00834 |
|  | Worst | 0.00832 | 35.20218 | 5.36624 | 29.04075 | 2.33717 | 33.64761 |
|  | Mean | 0.00832 | 2.31602 | 1.64951 | 4.58963 | 0.79998 | 3.49864 |
|  | STD | 6.99205e-7 | 6.64761 | 0.95211 | 7.00519 | 0.57348 | 7.02666 |
|  | Time | 16.87595 | 26.18215 | 59.46469 | 27.67845 | 72.13108 | 50.70354 |
| 10 | Best | 0.00831 | 0.00831 | 0.07548 | 0.00831 | 0.06963 | 0.00831 |
|  | Worst | 0.00831 | 32.91613 | 1.68895 | 2.29155 | 1.00556 | 0.06886 |
|  | Mean | 0.00831 | 2.76042 | 0.67766 | 0.09047 | 0.46844 | 0.01033 |
|  | STD | 1.86390e-9 | 8.19100 | 0.39728 | 0.41612 | 0.22787 | 0.01106 |
|  | Time | 33.51828 | 142.78045 | 257.42061 | 38.66741 | 167.58195 | 125.86006 |
| 15 | Best | 0.00831 | 0.00831 | 0.07948 | 0.00831 | 0.08545 | 0.00831 |
|  | Worst | 0.00831 | 12.14883 | 1.15398 | 0.06887 | 0.52837 | 0.00831 |
|  | Mean | 0.00831 | 0.59953 | 0.41119 | 0.01033 | 0.29428 | 0.00831 |
|  | STD | 2.46486e-12 | 2.25446 | 0.25924 | 0.01105 | 0.11665 | 2.73934e-13 |
|  | Time | 39.93289 | 67.50158 | 467.88354 | 39.44015 | 156.17842 | 90.21326 |
| 20 | Best | 0.00831 | 0.00831 | 0.08876 | 0.00831 | 0.07060 | 0.00831 |
|  | Worst | 0.00831 | 2.92471 | 0.86462 | 0.06887 | 0.21607 | 0.00831 |
|  | Mean | 0.00831 | 0.38322 | 0.32842 | 0.01033 | 0.14527 | 0.00831 |
|  | STD | 8.02622e-13 | 0.89581 | 0.22280 | 0.01106 | 0.03847 | 6.01556e-13 |
|  | Time | 69.00338 | 108.57437 | 996.30429 | 61.83259 | 234.38249 | 127.64768 |

**Table SI.14:** Estimated parameter values for the LK chemotherapy model with *N*_*p*_ = 5, 10, 15, and 20. Recall the true values are *V* (0) = 100, *ρ* = 0.1, *e* = 0.06, and *k* = 0.5 as listed in Table 4.

| $N_p$ | Parameter | | TSA | PSO | FA | DE | ABC | TLBO |
| --- | --- | --- | --- | --- | --- | --- | --- | --- |
| 5 | $V(0)$ | Best | 101.78688 | 98.42732 | 170.61801 | 99.24084 | 110.73609 | 101.95683 |
|  |  | Worst | 101.53931 | 534.48721 | 609.99878 | 0.00160 | 215.53352 | 1000.0 |
| | $\rho$ | Best | 0.09543 | 0.07284 | 0.07237 | 0.10394 | 0.05869 | 0.09495 |
|  |  | Worst | 0.09536 | 10.0 | 0.30223 | 2.37069 | 0.22726 | 3.81647 |
| | $e$ | Best | 0.06173 | 0.06152 | 1.05928 | 0.05421 | 0.64645 | 0.09495 |
|  |  | Worst | 0.06152 | 10.0 | 4.24629 | 4.54400 | 3.08165 | 9.99045 |
| | $k$ | Best | 0.54367 | 0.76049 | 9.82414 | 0.43252 | 10.0 | 0.06255 |
|  |  | Worst | 0.54272 | 10.0 | 10.0 | 6.20443 | 10.0 | 10.0 |
| 10 | $V(0)$ | Best | 101.78698 | 101.78707 | 98.41183 | 101.78695 | 106.05849 | 101.78695 |
|  |  | Worst | 101.79283 | 1000.0 | 181.09906 | 1000.0 | 143.95651 | 98.33367 |
| | $\rho$ | Best | 0.09543 | 0.09543 | 0.08178 | 0.09543 | 0.12432 | 0.09542 |
|  |  | Worst | 0.09543 | 1.45716 | 0.13181 | 0.19291 | 0.17382 | 0.07249 |
| | $e$ | Best | 0.06173 | 0.06173 | 0.42172 | 0.06173 | 0.04541 | 0.06173 |
|  |  | Worst | 0.06175 | 10.0 | 1.50469 | 0.16548 | 2.25252 | 0.78518 |
| | $k$ | Best | 0.54367 | 0.54367 | 4.64403 | 0.54367 | 0.27828 | 0.54367 |
|  |  | Worst | 0.54384 | 10.0 | 10.0 | 0.52852 | 10.0 | 10.0 |
| 15 | $V(0)$ | Best | 101.78696 | 101.78688 | 110.83152 | 101.78698 | 132.79364 | 101.78700 |
|  |  | Worst | 101.78702 | 685.69583 | 67.76703 | 98.33368 | 115.58346 | 101.78696 |
| | $\rho$ | Best | 0.09542 | 0.09543 | 0.07539 | 0.09543 | 0.08677 | 0.09543 |
|  |  | Worst | 0.09543 | 10.0 | 0.11822 | 0.07249 | 0.14496 | 0.09543 |
| | $e$ | Best | 0.06173 | 0.06173 | 0.67521 | 0.06173 | 0.13985 | 0.06173 |
|  |  | Worst | 0.06173 | 0.67277 | 0.36377 | 0.78518 | 1.32613 | 0.06173 |
| | $k$ | Best | 0.54367 | 0.54367 | 7.81284 | 0.54367 | 1.24506 | 0.54367 |
|  |  | Worst | 0.54367 | 0.04378 | 3.64396 | 10.0 | 7.47071 | 0.54367 |
| 20 | $V(0)$ | Best | 101.78698 | 101.78697 | 86.14816 | 101.78693 | 94.48461 | 101.78698 |
|  |  | Worst | 101.78697 | 1000.0 | 92.29330 | 98.33368 | 107.58939 | 101.78717 |
| | $\rho$ | Best | 0.09543 | 0.09543 | 0.09626 | 0.09543 | 0.07435 | 0.09543 |
|  |  | Worst | 0.09543 | 0.13395 | 0.16560 | 0.07249 | 0.11323 | 0.09543 |
| | $e$ | Best | 0.06173 | 0.06173 | 0.13664 | 0.06173 | 0.62648 | 0.06173 |
|  |  | Worst | 0.06173 | 2.36824 | 1.37571 | 0.78518 | 1.34899 | 0.06173 |
| | $k$ | Best | 0.54367 | 0.54367 | 1.28479 | 0.54367 | 7.93727 | 0.54367 |
|  |  | Worst | 0.54367 | 10.0 | 6.94029 | 10.0 | 10.0 | 0.54367 |

